# Reduced levels of inositol hexakisphosphate kinase (IP6K) impair life-cycle transitions and the intracellular development of *Trypanosoma cruzi* within human cardiomyocytes

**DOI:** 10.64898/2026.01.21.700787

**Authors:** Bryan E. Abuchery, Thaise L. Teixeira, Vitor L. da Silva, Rauni B. Marques, Fernanda M. Gerolamo, Anuj Shukla, Carolina M. C. Catta-Preta, Bruno A. Santarossa, Craig Lapsley, Yete G. Ferri, Maria Cristina M. Motta, Samuel C. Teixeira, Miguel A. Chiurrilo, Noelia M. Lander, Simone G. Calderano, Eduardo M. Reis, Henning J. Jessen, Richard McCulloch, Marcelo S. da Silva

## Abstract

*Trypanosoma cruzi* is the etiological agent of Chagas disease. During its life cycle, *T. cruzi* undergoes several key differentiation processes that are essential for its survival. The precise mechanisms that regulate these processes remain elusive, and any interference in this cycle would represent a breakthrough in the development of effective therapy against Chagas disease. Here, after depleting a single *IP6K* allele of *T. cruzi*, we observed that key differentiation processes (metacyclogenesis, amastigogenesis and trypomastigogenesis) were profoundly impaired. Epimastigote forms of IP6K-deficient *T. cruzi* exhibited morphological alterations and reduced metacyclogenesis. IP6K-deficient metacyclic forms had reduced infective potential in human cardiomyocytes. IP6K-deficient amastigote forms showed impaired ability to transform into trypomastigotes, with most of the population egressing from human cardiomyocytes without completing trypomastigogenesis. Together, our results suggest that IP6K is critical to sustain the *T. cruzi* life cycle. Since disruption of both *IP6K* alleles was lethal and the primary structure of IP6K shares only ∼25% similarity with its human homolog, this kinase emerges as a promising target for drug development against Chagas disease.

**Graphical Abstract:** 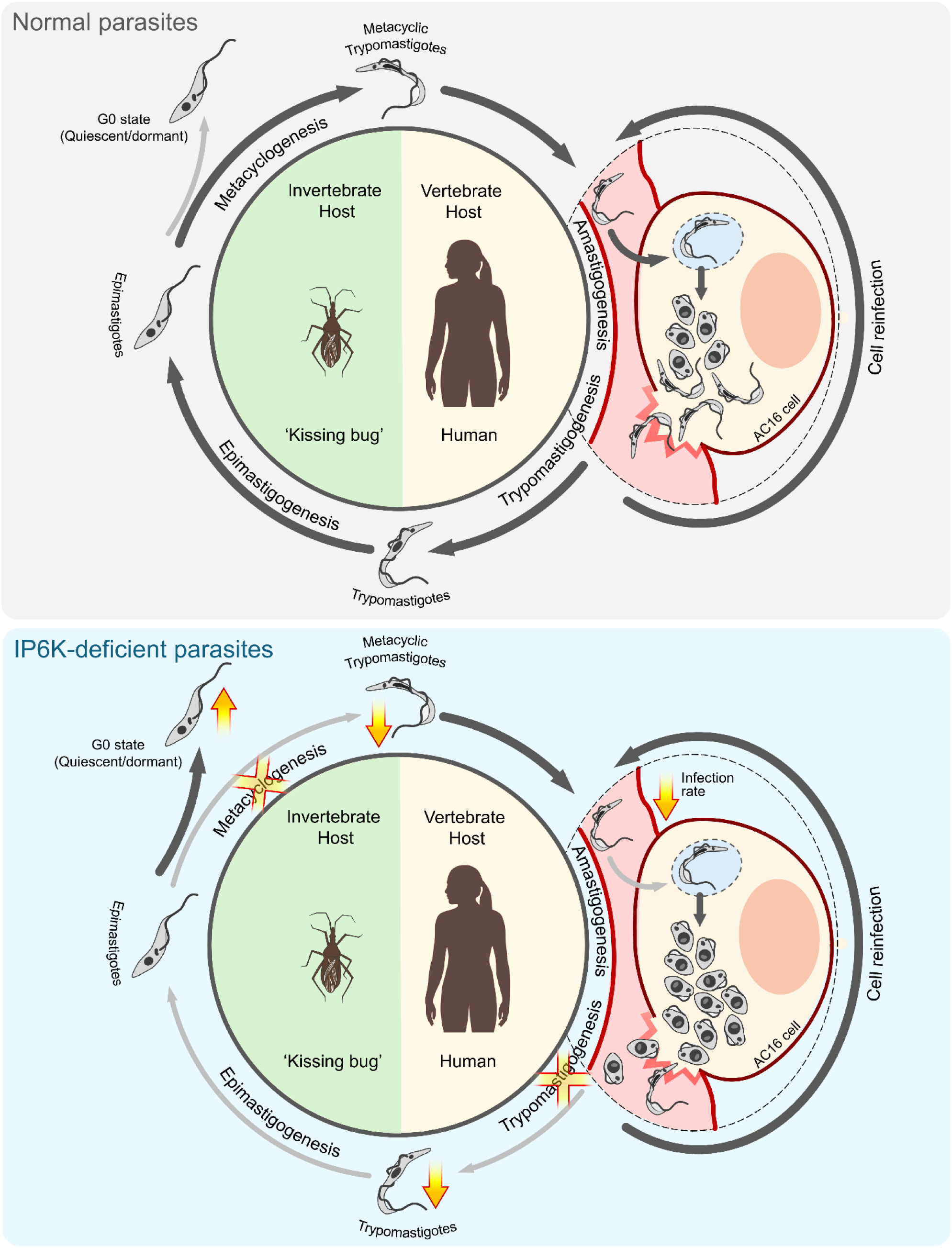

## Introduction

American trypanosomiasis, also known as Chagas’ disease (CD), is a neglected tropical disease that affects 6–7 million people worldwide, mainly in Latin America (Khare et al., 2016; Browne et al., 2017). CD is vector-borne, and its etiological agent is the single-celled eukaryote *Trypanosoma cruzi*, a protozoan parasite that, although it can infect and multiply in various organs, presents a preferential tropism for human cardiac tissue (reviewed in Rassi et al., 2000). Clinical forms of human CD are usually divided into two phases: acute and chronic (Rassi et al., 2010; Martín-Escolano et al., 2022). The acute phase occurs 4 to 8 weeks after infection and is usually asymptomatic, though it can be severe in children and can lead to death in up to 5% of diagnosed cases (Lewis et al., 2014). In general, most infected individuals eliminate the parasites from the bloodstream during this period (4 – 8 weeks) and then may enter the chronic phase, which is characterized by the absence of clinical manifestations for many years. After this long period, by a still unexplained mechanism, possibly related to parasite dormancy (quiescence) (Ward et al., 2020), clinical manifestations begin to appear, such as chronic Chagas cardiomyopathy (in 20–30% of infected individuals) and digestive megasyndromes (in ∼ 10% of cases) (Köberle, 1970; Arantes et al., 2004; Martín-Escolano et al., 2022). There are currently no effective vaccines against CD, and the first-line treatments rely on drugs developed several decades ago, which are associated with significant side effects (Urbina and Docampo, 2003; Naula et al., 2005; Khare et al., 2016; Browne et al., 2017).

The traditional view of the *T. cruzi* life cycle comprises two hosts: an invertebrate (usually a triatomine insect) and a vertebrate (usually a mammal), and four developmental forms: metacyclic trypomastigotes, amastigotes, bloodstream trypomastigotes and epimastigotes (Martín-Escolano et al., 2022). Human infection is usually initiated when urine/feces of the triatomine insect vector, contaminated with metacyclic trypomastigote forms of *T. cruzi*, comes into contact with the bite wound, allowing access to the bloodstream. Once in the blood, metacyclics can invade virtually any cell type. Within a cell, metacyclics escape the parasitophorous vacuole and initiate the transition to amastigote forms in a process called amastigogenesis. The amastigotes start multiplying within the host cell until they initiate a transition to non-proliferative trypomastigote forms (trypomastigogenesis). Next, the trypomastigotes egress the host cell, reaching the bloodstream, where they can reinfect other cells or be ingested by another triatomine during the blood meal. Once in the digestive tract of the insect, trypomastigotes transform into the proliferative epimastigote form (epimastigogenesis) that can migrate to the hindgut/rectum and differentiate into metacyclic trypomastigotes, a process named metacyclogenesis. *T. cruzi* metacyclic forms are then released in the feces/urine, enabling the life cycle to be completed (Martín-Escolano et al., 2022).

The transitions (cell differentiation) among *T. cruzi* developmental forms are triggered by environmental factors (e.g. starvation and changes in pH/temperature) and are regulated by protein kinases/phosphatases that control the phosphorylation/dephosphorylation of specific proteins responsible for sustaining its life cycle (Belaunzarán et al. 2009, Chiurillo et al., 2021, Marcelino et al., 2023). Thus, pathways involving kinases/phosphatases that may impair the standard progression of the *T. cruzi* life-cycle transitions are potential intervention points for an effective treatment against CD.

A signaling pathway that encompasses several kinases and seems to be of paramount importance for the *T. cruzi* life cycle is the inositol phosphates (IPs) pathway (Mantilla et al., 2021). The IPs pathway is a complex and dynamic route that involves a series of steps for the synthesis of inositol phosphates and inositol pyrophosphates (PP-IPs). The downstream steps of the IPs pathway culminate in the formation of inositol hexakisphosphate (IP_6_), the main substrate of inositol hexakisphosphate kinase (IP6K). The canonical function of IP6K is to synthesize the inositol pyrophosphate IP_7_ by transferring a β-phosphate group to position 1 or 5 of IP_6_ (Chakkour and Greenberg, 2024), leading to the formation of 1-IP_7_ or 5-IP_7_ (PP-IPs), water-soluble molecules involved in a wide range of cellular processes, including homologous recombination (Luo et al., 2002; Jadav et al., 2013), regulation of telomere homeostasis (Saiardi et al., 2005; Banfic et al., 2016), and regulation of fungal virulence (Lev et al., 2015; Desmarini et al., 2020).

Curiously, a growing body of studies has identified non-canonical functions beyond their catalytic role in PP-IPs synthesis. In mammals, three IP6K isoforms have been identified (IP6K1, IP6K2, and IP6K3) and among them, IP6K1 appears to play particularly diverse roles. IP6K1 contributes to neuronal migration through direct interactions with α-actinin and Arp2 (Fu el al., 2017; Qi et al., 2023). Moreover, IP6K1 also associates with the translation initiation complex on ribosomes and interacts with the mRNA decapping complex, promoting P-body assembly and regulating the abundance of decapping proteins EDC4, DCP1A/B, and DCP2 through mechanisms that remain to be elucidated (Shah and Bhandari, 2021). Also, IP6K1 has been implicated in promoting protein O-GlcNAcylation in human hepatocytes by an undefined pathway (Mukherjee et al., 2021). Thus, these findings highlight the emerging view that IP6Ks perform roles independent of their canonical catalytic activity in PP-IP synthesis.

Trypanosomatids encode only one isoform of IP6K and most studies involving this kinase are associated with its catalytic activity in the synthesis of PP-IPs. Mantilla et al. (2021) observed that perturbation of the IP pathway through the depletion of IPMK, IP5K and IP6K caused mild harmful effects in *T. cruzi* epimastigotes but significantly impaired the virulence of *T. cruzi* metacyclic trypomastigotes in Vero cells, suggesting that highly phosphorylated IPs are important for the maintenance of the *T. cruzi* life cycle. Interestingly, the authors were able to generate a lineage with only a single IP6K allele deleted, and the levels of PP-IPs in this lineage showed no significant differences, suggesting that IP6K may be essential for this organism due to roles unrelated to PP-IP synthesis. Besides, Cordeiro et al. (2017) showed that the PP-IPs pathway in *Trypanosoma brucei* is linked to IPs synthesis in acidocalcisomes. Furthermore, Cestari and Stuart (2015) observed that specific steps in the IPs pathway, possibly related to IP6K, regulate antigenic switching in *T. brucei* by controlling telomere homeostasis via shelterin components. These diverse phenotypes in *T. brucei* suggest that unexplored functions may have been overlooked for *T. cruzi* IP6K.

Here, we demonstrate that IP6K is partially conserved across eukaryotes, including the Euglenozoa group, which encompasses *T. cruzi*. *In silico* structure modeling predicts that *T. cruzi* IP6K retains its catalytic domain (IPK domain), followed by large intrinsically disordered regions. Using CRISPR-Cas9, we investigated the impact of a single-allele depletion of *IP6K* on the life cycle and on the main developmental forms of *T. cruzi*. We observed that IP6K-deficient *T. cruzi* epimastigote forms showed morphological alterations, impaired proliferation, and compromised metacyclogenesis. However, the PP-IPs levels (1-IP_7_, 5-IP_7_, and 1,5-IP_8_) did not undergo significant alterations. Moreover, IP6K-deficient epimastigotes displayed a portion of the population arrested in the G0 state/G1 phase (G0/G1) without significant DNA damage, suggesting an increase in the proportion of cells not committed to the cell cycle. In addition, *T. cruzi* IP6K-deficient metacyclic trypomastigotes exhibited reduced infectivity (invasion rate) in human cardiomyocytes, and the few IP6K-deficient amastigotes that were able to survive and proliferate after infection showed impaired trypomastigogenesis within human cardiomyocytes. Interestingly, most of the IP6K-deficient population egressed from human cardiomyocytes as amastigotes, without completing trypomastigogenesis. Furthermore, IP6K-deficient amastigotes displayed a high replication rate, suggesting a dysregulated proliferation. Transcriptomics analysis revealed that the mRNA expression levels of MASPs, trans-sialidases, and mucins in epimastigote and trypomastigote forms were altered in response to IP6K deletion, which may explain the impaired life cycle transitions observed. Taken together, our findings demonstrate that IP6K, possibly independently of PP-IPs, is fundamental to sustaining the life cycle of *T. cruzi*.

## Materials and Methods

### Phylogenetic analysis

The IP6K amino acid sequences (TcCLB.504213.90 – Esmeraldo-Like and TcCLB.506985.60 – Non-Esmeraldo-Like) were downloaded from TriTrypDB (tritrypdb.org/tritrypdb/app) and used to blast for orthologous sequences both in TritrypDB and Protein Data Bank (PDB). Orthologous amino acid sequences were put in one txt file, cleaned and dereplicated using specific packages in R. The file was used to generate a mapping file consisting of sequence identifiers by organism name and phylum. The sequences were aligned using Clustal Omega (Sievers et al., 2011) due to size of the fasta file using the command-line program and trimmed to remove misaligned columns using trimAl v1.4.rev15 (Capella-Gutiérrez et al., 2009) at a 0.1 gap threshold. To infer a phylogenetic tree, the RAxML-NG (Randomized Axelerated Maximum Likelihood – Next Generation) program was used. RAxML-NG infers Maximum Likelihood based on iteratively performing integrated series of Subtree Pruning and Regrafting (SPR) events, which allows it to quickly for the best-fit ML tree (Kozlov et al., 2019) using base frequencies provided by the best fitting model (Le and Gascuel, 2008).

Briefly, the Multiple Sequence Alignment (MSA) is read to ensure it does not contain sites of undetermined characters, sequences with undefined characters, duplicate taxon names or any other format errors (Kozlov et al., 2019). The best-fit model for protein phylogenetic tree construction, according to the Bayesian Information Criterion (BIC) is determined using IQ-Tree (Nguyen et. al., 2015). A tree is then inferred the determined best-fit model [LG+I+G4; where LG is the model with invariable sites (+I) and gamma rate heterogeneity (+G4)]. Default parameters performed 20 tree searches using 10 random and 10 parsimony-based starting trees. The best-scoring topology tree was picked and then a standard non-parametric bootstrapping by re-sampling alignment columns and re-inferring the tree for each bootstrap replicate (1000 replicates) of the MSA was done to get accurate support values (Kozlov et al., 2019).

By default, RAxML-NG performs MRE-based bootstrapping test after every 50 replicates and terminates once the diagnostic statistics drops below the computational cutoff value (Kozlov et al., 2019). The default cutoff value for the MRE-based bootstrapping in RAXML-Ng is typically set at a statistical level of 0.03 (or 3%). BS (bootstrap) support values are then mapped onto the best-scoring/best-known ML tree, and a Tree Likelihood Evaluation is finally done to compute the likelihood of a given fixed tree topology by optimizing model and/or branch length parameters on that fixed tree (Kozlov et al., 2019).

The run was performed in the University of São Paulo system grid server, (AMD EPYC 7643 48-Core Processor, 96 cores, 1007 GB RAM) consisting of parallelization scheme autoconfig: 4 worker(s) x 128 thread(s) that took 194882.035 seconds to run. In this case, the diagnostic statistics dropped below the computational cutoff value at 600 bootstraps hence number of bootstraps was 600. The tree was visualized through Chiplot Online (Xie et al., 2023). To construct the Euglenozoa phylogenetic trees, we used the same method described above (with tree building completing at 150 bootstraps and 155.350 seconds run time). In this case, orthologous sequences occurring only in TritrypDB were used. The similarity matrix used to prepare heatmap visualization file was obtained from the Clustal-Omega alignment result files and the tree visualized through Interactive Tree of Life v5 (Letunic and Bork, 2021).

### T. cruzi IP6K structures prediction and molecular docking

To compare the primary and secondary structures of *T. cruzi* IP6K and its human isoforms, we firstly used the amino acids sequence (primary structure) of IP6K from Esmeraldo-like haplotype (TcCLB.504213.90) to blast (using the blastp program) for human orthologs from the NCBI database (https://blast.ncbi.nlm.nih.gov/Blast.cgi). These sequences were then put in one fasta file and aligned using an online ClustalOmega tool (https://www.ebi.ac.uk/jdispatcher/msa/clustalo). The alignment was visualized by Jalview V 2.11.5.0. The amino acid alignment was then colored using Jalview’s amino acid percentage similarity color scheme. Higher percentage similarity results in a more intense color, typically a deeper blue, while lower similarity leads to a lighter blue or white. The mapping of *T. cruzi* IP6K secondary structures on the alignment was predicted by EMBL-EBI’s Pictorial database of 3D structures in the Protein Data Bank (http://www.ebi.ac.uk/thornton-srv/databases/pdbsum). The coordinates of the highlighted structures in black boxes were estimated using *E. histolytica* IP6K’s known structures as reference.

For the molecular docking and tertiary structure prediction analysis, the amino acids sequence of *T. cruzi* IP6K (TcCLB.504213.90) was submitted to AlphaFold3 (Abrahamson et al., 2004). To predict which amino acids are important during IP_7_ synthesis (from IP_6_), we added in the simulations an ATP (adenosine triphosphate) molecule and two Mg^2+^ ions (Wang et al., 2014, Mantilla et al., 2021). Autodock Vina, v1.2.7 (Trott and Olson, 2010) was used to predict the ligand binding site and thus the interaction of *T. cruzi* IP6K + ATP + 2 Mg^+^ + IP_6_. The tridimensional structure of the IP_6_ ligand (DrugBank ID DB14981) used was derived from RCSB Protein Data Bank and used in our analyses. The best conformation with the best fit predicted an affinity (kcal/mol) energy of - 5.6. The results were visualized using ChimeraX V1.9 (https://www.rbvi.ucsf.edu/chimerax/) and BioDiscovery Viewer (https://discover.3ds.com/discovery-studio-visualizer-download).

### Cellular cultures and maintenance

*T. cruzi* epimastigote forms (CL Brener strain) were cultured at 27 °C in liver-infusion tryptose (LIT) medium supplemented with 10% fetal bovine serum (FBS) (Cultilab) and 1% (v/v) antibiotic solution (streptomycin/penicillin).

*T. cruzi* metacyclic forms were obtained by inducing metacyclogenesis from epimastigote forms using Grace medium (Gibco). The metacyclic forms were purified on a DEAE cellulose column according to Teixeira and Yoshida (1986).

Tissue-culture cell-derived trypomastigote forms (TCTs) were obtained by infecting LLCMK2 cells with purified *T. cruzi* metacyclic forms (MOI 10:1). Usually, after 120 h of invasion, *T. cruzi* TCTs egress from the LLCMK2 cells and are collected by centrifugation (1000 *g*, 7 min).

Of note, the different life forms of *T. cruzi* lineages containing the T7/Cas9 construct (parental control) and *IP6K*^-/+^ were obtained and maintained in the same conditions but supplemented with the respective antibiotics: 100 μg.mL^-1^ G418 sulfate for Control and 10 μg.mL^-1^ blasticidin for *IP6K*^-/+^.

Immortalized LLCMK2 cells and AC16 human cardiomyocytes were maintained respectively in DMEM (Cytiva) and DMEM F12 (Gibco) medium, both supplemented with 10% FBS at 37 °C. LLCMK2 cells were used only to maintain the *T. cruzi* TCTs forms and AC16 human cardiomyocytes were used in the assays involving host-pathogen interaction. LLC-MK2 cells were obtained from BCRJ (#code 0146) (https://bcrj.org.br/celula/llc-mk2-original/) and AC16 cells were acquired from Merck Millipore (#SCC109).

### Growth curves

For the growth curves, *T. cruzi* lineages were started at a concentration of 2×10^6^ cells.mL^-1^ and counted daily for 8 days. The counting was done using a Neubauer chamber. The final concentration was given by the following formula: <u>*x*</u> *quadrants.dilution*^−1^. 10^4^. The standard deviation (SD) in each growth curve was estimated from three independent experiments.

### Generation of T. cruzi IP6K mutants using the CRISPR-Cas9 system

The *T. cruzi* (CL Brener) genome editing using the CRISPR-Cas9 approach was based on the system developed by Costa et al. (2018) and Beneke et al. (2017) (Beneke et al., 2017; Costa et al., 2018). CL Brener is a naturally occurring hybrid *T. cruzi* strain of distinct lineages (Esmeraldo-like and Non-Esmeraldo-like) (El-Sayed et al., 2005). Due to that, the CRISPR-Cas9 approach requires extra attention. On the other hand, unlike mammals, trypanosomatids do not present IP6K paralogs (Cordeiro et al., 2017; Mantilla et al., 2021). Also, in *T. cruzi* CL Brener, *IP6K* is a single-copy gene located on chromosome 11, as confirmed by whole-genome sequencing analyses (Supplementary Figure 2A) and local alignment searches in the integrated database TriTrypDB (https://tritrypdb.org), which facilitates genome editing using the CRISPR/Cas9 approach.

Briefly, the DNA donors (harboring the resistance genes against blasticidin and puromycin) were amplified using Phusion High-Fidelity DNA polymerase (Thermo Fischer Scientific), specific primers (Supplementary Table 1 – Upstream foward and Downstream reverse primers), and template plasmids provided as a gift by Professor John Kelly (London School of Hygiene & Tropical Medicine, UK). The amplification of the DNA responsible for sgRNA transcription (‘sgRNA’), which enabled the cleavage of both *IP6K* alleles in the 5’ and 3’ regions, was carried out through a template-free PCR reaction using specific primers common to the Esmeraldo-like and Non-Esmeraldo-like haplotypes (Supplementary Table 1 – Upstream and Downstream ‘sgRNA’ primers). In this reaction, it was used a Phusion High-Fidelity DNA polymerase (Thermo Fischer Scientific) in a final volume of 20 μL. The “Eukaryotic Pathogen CRISPR guide RNA/DNA Design Tool” (http://grna.ctegd.uga.edu/) was used to define the 20-base pair sequence (protospacer) of the ‘sgRNA’. This tool generates several specific sequences that have the PAM motif (5’-NGG-3’) for the region to be cleaved, eliminating the possible sequences that would cause unspecific breaks (off-targets). Prior to transfection, the DNA donors and ‘sgRNA’ PCR products were purified on silica columns (QIAquick PCR purification Kit – Qiagen) and eluted in 50 μL of TE (10 mM Tris, 1 mM EDTA) previously filtered on a 0.22 μm nylon membrane.

Two samples of exponentially growing *T. cruzi* T7/Cas9 epimastigotes (1×10^8^ total parasites) were used during transfection. These samples were centrifuged, washed in PBS (137 mM NaCl, 2.7 mM KCl, 10 mM Na_2_HPO_4_ and 2 mM KH_2_PO_4_, pH 7.4) and resuspended in 200 µL transfection buffer (120 mM KCl, 0.15 mM CaCl_2_, 2 mM EDTA, 10 mM K_2_HPO_4_/KH_2_PO_4_, 25 mM HEPES, 5 mM MgCl_2_).

Then, the resuspended samples were transferred to a 0.2 cm cuvette together with the purified PCR products or TE buffer (for control). The electroporation was carried out using the Nucleofector 2b (Lonza) according to the manufacturer’s instructions. Two pulses were applied using the X-014 program. Then, each sample was placed in 5 mL of LIT medium and incubated at 27 °C for 24 h. The day after electroporation, the *IP6K* KO sample was split into 3 flasks and, subsequently, *IP6K* KO samples and the control received the needed selection antibiotics (puromycin, blasticidin and/or G418 sulfate). The parasites were incubated at 27 °C in the presence of 5% CO_2_. The selection of the polyclonal population was achieved ∼14 days after transfection, when no live cells were found in the control group. The monoclonal population was obtained from the polyclonal population after limiting dilution. Of note, the attempt to obtain the *IP6K* double-null cells (*IP6K*^-/-^) was carried out in the same way as was described but using the IP6K^-/+^ cells as a background. Unfortunately, we were not successful in obtaining *IP6K* double-null cells.

### Genomic DNA (gDNA) extraction and PCRs

From each lineage analyzed, around 1×10^8^ *T. cruzi epimastigotes* (in exponential phase) were collected by centrifugation (1600 *g* for 5 min), washed twice in PBS, re-suspended in 4 mL of Lysis solution (200 mM Tris, 10 mM EDTA-pH 8.0, 50 μg.mL^-1^ proteinase K, and 0.5% SDS) and incubated overnight at 37 °C. Then, 2.1 mL Phenol (Sigma-Aldrich) was added, and the solution was gently mixed. Next, 2.1 mL of Chloroform (Merck) was added into the solution, which was then gently mixed and centrifuged (1600 *g* for 10 min). The aqueous phase (top phase) was collected and transferred to a clean tube. 10 μL RNase A (New England Biolabs) – from the stock 20 mg.mL^-1^ – was added to the aqueous phase and incubated at 37 °C for 30 min. Phenol/chloroform/isoamyl alcohol (25:24:1) v/v was added and the solution was mixed gently. The solution was centrifuged again and the aqueous phase collected into another tube. The gDNA was precipitated by the addition of 30 mM Sodium acetate and twice volumes of 100% ethanol (Merck). The solution containing gDNA was gently mixed and incubated at -20 °C for 30 min. The gDNA was collected, washed using 70% ethanol (v/v) and solubilized into 1 mL TE solution (10 mM Tris, 1 mM EDTA, pH 8.0).

The gDNA samples were stored at 4 °C until use. Typically, they were used in PCR reactions to confirm the generated strains. For this purpose, we used the primers previously designed (Supplementary Table 1), dNTPs (Thermo Fischer Scientific), and Platinum II *Taq* Hot-Start DNA polymerase (Invitrogen). The reactions were carried out using a T100 Thermal Cycler (BioRad).

For the whole genome sequencing (WGS), gDNA was extracted from each sample using the DNeasy Blood & Tissue Kit (Qiagen) according to the manufacturer’s instructions. The quality and integrity of the DNA samples were measured using Qubit Fluorometer 3.0 (Thermo Fischer Scientific).

### Reverse Transcription quantitative Polymerase Chain Reaction (RT-qPCR)

Total RNA from each *T. cruzi* lineage analyzed was isolated from 1×10^9^ epimastigotes (total cells) in exponential phase using TRIzol (Thermo Fisher Scientific) according to the manufacturer’s instructions. RNA integrity was checked by gel electrophoresis and through Qubit Fluorometer 3.0 (Thermo Fisher Scientific). Two micrograms of total RNA from each individual sample were used for cDNA synthesis using the iScript cDNA Synthesis Kit (BioRad), according to manufacturer’s instructions and the previously described protocol (de Waal et al. 2008). The RT-qPCR assays were carried out using the RealQ Plus 2x Master Mix Green (Ampliqon), based on the protocol previously described (de Castro-Assis et al., 2018) with slight modifications. All RT-qPCRs were performed in 10 µL reactions and the cycle threshold values (Cts) were obtained in a Real-Time PCR system (Applied Biosystems) using default settings. The relative gene expression data were obtained using the 2^-ΔΔCT^ method as previously described (Livak and Schmittgen, 2001). Due to its constitutive expression among all groups analyzed, glyceraldehyde-3-phosphate dehydrogenase (GAPDH) mRNA levels were used as endogenous control. Samples from the control group were normalized to an arbitrary value of 1. The data was expressed as means of fold change over control (normalized expression). Supplementary Table 2 contains all the primers used in our RT-qPCR analyses.

### Whole genome sequencing (WGS)

For the WGS, each gDNA sample was diluted to a final concentration of 50 ng.mL^-1^ (60 μL total) and sent to the company Plasmidsaurus Inc. (plasmidsaurus.com). The WGS was performed as specified: diploid eukaryote, 55 Mb genome with 30x coverage, Hybrid Oxford Nanopore Technology (ONT) + Illumina WGS, ONT data target 1.67 Gb per sample (total ONT data target = 5 Gb), Illumina data target 1.67 Gb per sample (total Illumina data target = 5 Gb).

### Whole genome off-target deletion analysis

ONT + Illumina paired-end reads from WT, T7/Cas9 (Control), and IP6K^-/+^ were aligned with BWA-MEM (v0.7.17) to chromosome-scale pseudomolecules. Pseudomolecules were generated by anchoring assembly contigs to the *T. cruzi* CL Brener Esmeraldo-like reference (TriTrypDB-68) using minimap2 (preset asm5), ordering/orienting placements into scaffolds recorded in an AGP. Analyses were restricted to assembled AGP “W” components. For copy-number profiling, the genome was tiled into non-overlapping 20 kb windows. Mean coverage per window was computed from sorted/indexed BAMs; log_2_(sample/WT) was calculated per window. Windows with near-zero WT depth were flagged and excluded from summaries. To correct library-wide depth differences, each sample’s log2 track was iteratively re-centered by subtracting a MAD-trimmed median until convergence.

Deletions were called on the re-centered 20 kb tracks under two criteria: strict, ≥ 5 consecutive windows with log_2_ ≤ −1.0; and mild, ≥ 3 consecutive windows with log_2_ ≤ −0.7. As IP6K^-/+^ derives from T7/Cas9 (Control), a background mask was defined as the union of T7/Cas9 (Control) strict+mild segments, expanded by one window (±20 kb). Overlapping IP6K^-/+^ segments were removed. Regions within ±40 kb of AGP joints were also excluded. Remaining intervals were annotated with mean re-centered log_2_ (IP6K^-/+^ and T7/Cas9 – Control), bin count, fraction of reads with mapping quality zero (MAPQ0), and split-read counts within ±2 kb of each interval edge. The differential amplitude Δlog_2_ was computed as the mean re-centered log_2_(IP6K^-/+^/WT) minus the mean re-centered log_2_(T7/Cas9-Control/WT), [equivalently, log_2_(IP6K^-/+^/T7/Cas9-Control)]. Final 20-kb candidates were required to satisfy all the following: ≥ 4 windows (≥ 80 kb), mean log_2_(IP6K^-/+^) ≤ −0.8, and Δlog_2_ ≤ −0.3. Bins with a MAPQ0 fraction ≥ 0.9 (i.e., < 10% uniquely mapped reads) were classified as low-mappability and excluded. Results were unchanged using a more permissive threshold of 0.8.

To assess smaller events, AGP “W” regions were tiled into 1 kb windows. Mean coverage was obtained with samtools bedcov (sum of per-base depth divided by window length), followed by the same log_2_ computation and re-centering. Small deletions were called using tiered criteria (single-bin log_2_ ≤ −1.5; runs of ≥ 2 bins at ≤ −1.2; runs of ≥ 3 bins at ≤ −1.0), merged, minus the T7/Cas9 (Control) background (merged calls, ±1 kb), and filtered to exclude ±10 kb around AGP joins. Final 1 kb candidates required bins ≥ 3, mean log_2_(IP6K^-/+^) ≤ −1.0, Δ ≤ −0.3, MAPQ0 < 0.9, and ≥ 2 split reads across the interval edges.

### Scanning electron microcopy analysis

For Scanning Electron Microscopy (SEM) analysis, epimastigote forms of *T. cruzi* lineages in exponential phase (2×10^7^ cells.mL^-1^) were collected and fixed in 200 μL of 2.5% glutaraldehyde (Sigma-Aldrich) diluted in 0.1 M Sodium cacodylate (Sigma-Aldrich) solution, pH 7.2 for 48 h. Next, aliquot from each sample (50 μL) were settled for adhering into 0.1% poli-L-lysine pre-treated microscope coverslips (Knittel Glaser). The samples were then incubated with 0.1 M Sodium cacodylate solution (pH 7.2), containing 1% Osmium tetroxide (Sigma-Aldrich) and 0.8% Potassium ferricyanide (Sigma-Aldrich) for 45 min. Next, the coverslips containing samples were washed in 0.1 M Sodium cacodylate buffer (pH 7.2) and dehydrated in a stepwise series of ethanol (Merck), followed by critical point drying in CO_2_ using a Leica EM CPD030 apparatus (Leica). The samples were sputtered with gold in a Balzers FL9496 unit (Postfach 1000 FL-9496 Balzers Liechtenstein) and observed/captured in an EVO 40 VPSEM scanning electron microscope (Zeiss).

### Flow cytometry analysis

For analysis of internal complexity and relative size of the cells, epimastigote forms of *T. cruzi* lineages in exponential phase (2×10^7^ cells.mL^-1^) were collected from the culture bottle and directly analyzed using CytoFLEX LX Flow Cytometer (Beckman Coulter). The internal complexity of the cell was measured based on the scattering signals detected at a 90° relative to the laser (side scatter – SSC) in area and height (A and H). The relative size of the cells was measured based on the scatter signal along the path of the laser (forward scatter – FSC) in area and height (A and H).

For the DNA content analyses, *T. cruzi* lineages in exponential phase (2×10^7^ cells.mL^-1^) were collected by centrifugation (1000 *g* for 7 min), washed twice using PBS, and fixed/permeabilized with 70% cold methanol overnight at 4 °C. The pellet containing cells was washed once with PBS and resuspended in 500 μL of PBS containing 10 μg.mL^-1^ PI (Propidium Iodide) or 0.5 μg.mL^-1^ DAPI (4′,6-diamidino-2-fenilindol) and 100 μg.mL^-1^ of RNase A. After 30 min of incubation at 37 °C, the cells were analyzed using the BD Accuri C6 Flow Cytometer (BD Biosciences) or the CytoFLEX LX Flow Cytometer (Beckman Coulter).

The FCS files generated were analyzed using FlowJo^TM^ v10 (BD – Becton, Dickinson & Company) or Floreada.io software (https://floreada.io/).

### Inositol polyphosphates analysis

From each *T. cruzi* lineage analyzed, around 1×10^9^ epimastigotes (in exponential phase) were collected by centrifugation (2000 *g* for 7 min), washed twice in BAG solution (116 mM NaCl, 5.4 mM KCl, 0.8 mM MgSO_4_, 5.5 mM *D*-Glucose, and 50 mM HEPES, pH 7.0), re-suspended in 1 M perchloric acid, sonicated with 40% amplitude (Sonics Vibra-Cell™) for 10 sec and kept on ice for 15 min (this sonication procedure was repeated three times). The samples were then centrifuged at 18000 *g* for 5 min at 4 °C and the supernatant of each sample was transferred to a new tube and kept on ice. The pellet of each sample was resuspended in 200 μL of RIPA buffer (50 mM Tris-HCl, pH 7.4, 150 mM NaCl, 1% NP40, 0.5% sodium deoxycholate, 0.1% SDS, and 1% protease inhibitor cocktail) and the total amount of protein was measured using BCA protein assay kit (Pierce™). The total amount of protein was used to normalize the amount of inositol polyphosphates present in each sample.

Next, TiO_2_ beads were prepared according to Wilson and Saiardi, (2018). Briefly, 4 mg TiO_2_ beads (Titansphere TiO 5 µm, GL Sciences Inc) were added into a new 1.5 mL tube and washed – using a vortex – with 1 mL ultrapure water. The water was removed after centrifugation (3500 *g* for 2 min) and this step was repeated but replacing water with 1 M HClO_4_. After the last step, the TiO_2_ beads were resuspended in 50 μL 1 M HClO_4_.

Then, the supernatant containing the cell lysates were added to the tube containing TiO_2_ beads and left rotating for 30 min at 4 °C. Then, the TiO_2_ beads were centrifuged at 3500 *g* for 2 min at 4 °C, the supernatant was removed and the beads were washed twice in 0.5 mL of cold 1 M HClO_4_. After carefully removing the HClO_4_, the inositol phosphates attached to the beads were eluted by addition of 200 μL 2.8% ammonium hydroxide following rotation for 5 min at 4 °C. The beads were precipitated by centrifugation at 3500 *g* for 2 min and the supernatant containing the inositol phosphates was collected and transferred to a new tube. This elution step was repeated one more time and the eluate was combined with the one collected previously, resulting in a total volume of ∼400 μL. Then, the eluate containing the inositol polyphosphates was lyophilized using a SpeedVac evaporator (Savant). The dried samples were then stored at −20 °C, until reconstitution and analysis by capillary electrophoresis-electrospray ionization mass spectrometry (CE–ESI–MS) (Qiu et al., 2023; Liu et al., 2025; Liu and Jessen, 2025) or polyacrylamide gel electrophoresis (PAGE) (Losito et al., 2009; Wilson et al., 2015).

The CE–ESI–MS analyses were carried out using a setup consisting of an Agilent 7100 CE system coupled to an Agilent 6495C Triple Quadrupole mass spectrometer equipped with an Agilent Jet Stream electrospray ionization source and a dedicated CE–MS interface. For selected measurements, the CE system was alternatively connected to a quadrupole time-of-flight (QToF) mass spectrometer using the same CE–MS adapter and sprayer kit. All measurements were performed in negative ionization mode. For the separations, we used bare fused-silica capillaries (100 cm length, 50 µm internal diameter). Before each set of measurements, the capillary was conditioned by flushing with 1 M NaOH for 10 min, followed by water for an additional 10 min. Depending on the type of analysis and the level of separation required, different background electrolytes (BGEs) were used. For general profiling of inositol polyphosphates, we used a BGE containing 35 mM ammonium acetate adjusted to pH 9.75 with ammonium hydroxide. For QqQ measurements, the sheath liquid composition was identical to that used for QToF measurements, except that no reference ions were added. The sheath liquid flow rate was maintained at 10 µL.min^-1^. Instrument control and data acquisition were carried out using Agilent MassHunter Workstation software (version 10.1). Detailed parameters are provided in Supplementary Table 3–4. For quantitative analysis, dried TiO_2_-enriched extracts were reconstituted in deionized water prior to CE–MS measurement. Absolute and relative quantification of inositol polyphosphates was performed using stable isotope-labeled internal standards that were spiked directly into each sample before analysis (Qiu et al., 2020). IP_5_, IP_6_, IP_7_, and IP_8_ species were quantified using defined amounts of their corresponding ^13^C_6_-labeled standards as internal references (Harmel et al., 2019). Isotopic standards were added at concentrations specified in Supplementary Table 5. These concentrations were selected to ensure accurate detection within the linear dynamic range of the instrument, allowing reliable and reproducible quantification across samples.

The PP-IPs analysis by PAGE was performed as described by Wilson et al. (2015). Briefly, samples were mixed (2:1) with Loading dye [0.05% (w/v) Orange G, 30% Glycerol, 1 mM EDTA] and resolved on 35% acrylamide/bisacrylamide (19:1) PAGE using TBE buffer (89 mM Tris-borate, 2 mM EDTA, pH of 8.3). 2 μL of 1 mM Phytic acid (IP_6_) (MedChemExpress) was used as a molecular standard. After electrophoresis, the gel was stained for 30 min, at RT, with Toluidine blue staining solution (20% methanol; 2% glycerol; 0.05% toluidine blue) and then destained with de-staining solution (20% methanol; 2% glycerol) for 2 h, changing the solution several times during this period. Pictures were taken after exposing the gel to a white light transilluminator.

### Immunofluorescence Assay (IFA) – FKN pattern analyses

Some cell cycle phases of *T. cruzi* are easily distinguishable by measuring the number of DNA-containing organelles (nucleus and kinetoplast) and flagellum. We used this approach to measure the different Flagellum-Kinetoplast-Nucleus (FKN) patterns in each *T. cruzi* sample analized. Each *T. cruzi* lineage was collected in exponential phase (2×10^7^ cells.mL^-1^) by centrifugation (1000 *g* for 7 min), washed twice with PBS, fixed for 10 min with 4% sterile paraformaldehyde (Merck) diluted in PBS, and added into a 0.1% poli-L-lysine pre-treated slide (Knittel Glaser). The slides were then permeabilized with 0.1% Triton X-100 (Sigma-Aldrich) (diluted in PBS) for 5 min and briefly washed with PBS. α-flagellum (primary antibody) developed in mouse (da Silva et al., 2013) (diluted 1:10 in 4% BSA solution) was added onto the slide, which was incubated overnight at 4 °C to minimize chances of non-specific interactions. Next, the slides were carefully washed with PBS and the secondary antibody (Alexa Fluor 555 anti-mouse IgG diluted 1:1000 in 4% BSA) was added, being incubated for 2 h at 4 °C protected from light. Then, the slides were washed carefully with PBS and left to dry out for few minutes. Vectashield containing DAPI (VectorLabs) was used as an anti-fade mounting solution and to stain nuclei and kinetoplast DNA. Images were acquired using the fluorescent microscope (Nikon Eclipse 80i) with 100x oil attached to a mercury lamp and a digital camera (Nikon DS-F1i).

DAPI-flagellum-stained *T. cruzi* epimastigotes were examined and the following FKN patterns were established: 2F2K2N (cytokinesis), 2F2K1N (G2/mitosis), 2F1K1N (late S/G2), 1F1K1N (G1/early-middle S), and others (aberrant cells, i.e., profiles that do not fall within the established criteria).

### EdU incorporation assays and ‘click’ chemistry reaction

Exponentially growing *T. cruzi* epimastigotes (2×10^7^ cells.mL^-1^) were incubated with 100 µM 5-ethynyl-2’-deoxyuridine (EdU) (Thermo Fischer Scientific) for different time periods (ranging from 1 h to 18 h according to the approach used) at 27 °C. The parasites were then harvested by centrifugation (1000 *g* for 7 min), washed twice in PBS, fixed for 10 min with 4% sterile paraformaldehyde (Merck) diluted in PBS, washed twice, and re-suspended in 50 µL of PBS. Then, 25 µL of the cell suspension was loaded onto 0.1% poly-L-lysine pre-treated slides (Knittel Glaser). The cells were permeabilized with 0.1% Triton X-100 (Sigma-Aldrich) (diluted in PBS) for 5 min and then briefly washed using PBS. To detect incorporated EdU, we used 50 µL of Click-iT EdU detection solution (3 mM CuSO_4_, 100 mM C_6_H_8_O_6_, and 5 mM Alexa fluor azide 488 diluted in PBS) for 1 h protected from light. For details about EdU procedure, see (Salic and Mitchison, 2008). Finally, the cells were washed twice with PBS. Vectashield containing DAPI (VectorLabs) was used as an anti-fade mounting solution and to stain nuclei and kinetoplast DNA. Images were acquired using the fluorescent microscope (Nikon Eclipse 80i) with 100x oil attached to a mercury lamp and a digital camera (Nikon DS-F1i). Images were further analyzed using ImageJ (version IJ 1.46r) (National Institutes of Health).

### Cell cycle analysis

The profile 2F2K2N (from the FKN approach) was used to estimate the percentage of *T. cruzi* epimastigote cells in cytokinesis (C). Using this parameter, we were able to estimate the duration of cytokinesis according to the Williams equation (Williams, 1971):

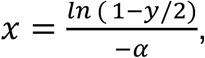

where x is the cumulative time within the cycle until the end of the stage in question, y is the cumulative % of cells up to and including the stage in question (expressed as a fraction of one unit), and α is the specific growth rate.

To estimate the G2 and Mitosis phases (G2+M) length, we added EdU in the medium containing exponential growing epimastigotes and collected samples every 15 min, proceeding with ‘click’ chemistry reaction, until a parasite containing two EdU-labeled nuclei (2N2K) was observed (this time corresponds to the length of G2+M phases).

To estimate the S-phase duration, we measured the proportion of EdU-labeled cells after 1 h EdU pulse. The S-phase duration was estimated according to the Stanners and Till equation (Stanners and Till, 1960):

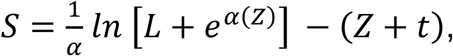

where *L* is the proportion of cells exhibiting EdU-labeled nuclei, *α* = *ln* 2.T^-1^ (*T* = doubling time, expressed in hours), Z = G2 + M + cytokinesis, and *t* is the duration of the EdU pulse in hours (h).

Finally, the duration of G1 phase was estimated by the difference between the doubling time (dt) and the sum of the other phases (S+G2+M+C). Of note, all these calculations were made with the help of the online software CeCyD, available at the address https://cecyd.vital.butantan.gov.br/ (da Silva et al., 2020). The doubling time was estimated by taking the values at exponential phase (growth cruves) and using the Doubling Time software (http://www.doubling-time.com).

### Immunofluorescence Assay (IFA) using α-γH2A

The presence of DNA damage can be monitored by signaling and activating important players involved in the response, such as γH2A (Glover and Horn, 2012). To verify the presence of DNA damage, we used an α-γH2A antibody (da Silva et al., 2019). As positive control [C(+)], the *T. cruzi* control lineage was incubated with 500 µg.mL^-1^ phleomycin (InvivoGen) for 2 h (Glover and Horn, 2012). As negative control [C(–)], the *T. cruzi* control lineage was processed replacing α-γH2A with PBS (v/v). *T. cruzi* lineages were collected by centrifugation (1000 *g* for 7 min), washed twice with PBS, fixed for 10 min with 4% sterile paraformaldehyde (Merck) diluted in PBS, and added into a 0.1% poli-L-lysine pre-treated slide (Knittel Glaser). The slides containing cells were processed as described in FKN pattern analyses, replacing α-flagellum (mouse) with α-γH2A (rabbit) antibody, which was diluted 1:500 in 4% BSA solution. As a secondary antibody we use Alexa Fluor 555 anti-rabbit IgG (Thermo Fischer Scientific) diluted 1:1000 in 4% BSA which was incubated for 2 h at 4 °C protected from light. Then, the slides were washed carefully with PBS and left to dry out for few minutes. Vectashield containing DAPI (VectorLabs) was used as an anti-fade mounting solution and to stain nuclei and kinetoplast DNA. Images were acquired using the fluorescent microscope (Nikon Eclipse 80i) with 100x oil attached to a mercury lamp and a digital camera (Nikon DS-F1i).

### Terminal deoxynucleotidyl transferase dUTP nick end labeling (TUNEL) assay

Exponentially growing *T. cruzi* epimastigotes (2×10^7^ cells.mL^-1^) were collected by centrifugation (1000 *g* for 7 min), washed twice with PBS, fixed for 10 min with 4% sterile paraformaldehyde (Merck) diluted in PBS, and added into a 0.1% poli-L-lysine pre-treated slide (Knittel Glaser). The DeadEnd™ Fluorometric TUNEL System kit (Promega) was used according to the manufacturer’s instructions to detect DNA fragmentation. The following controls were included: a negative control (C−), where the reaction was carried out without the rTdT enzyme; and a positive control (C+), where the fixed parasites were pre-treated with DNase I (ThermoFisher) for 10 min. At the end of the TUNEL reactions, the slides were washed carefully with PBS and left to dry out for few minutes. Vectashield containing DAPI (VectorLabs) was used as an anti-fade mounting solution and to stain nuclei and kinetoplast DNA. Images were acquired using the fluorescent microscope (Nikon Eclipse 80i) with 100x oil attached to a mercury lamp and a digital camera (Nikon DS-F1i).

### Analysis of cells not committed to the cell cycle (quiescence)

To quantify cells not committed to the cell cycle (quiescent cells), we used an EdU-negative selection. This approach consists of performing an EdU incorporation for a long period (longer than the sum of the G2+M+C+G1 cell cycle phases) and measuring the cells that could not uptake this thymidine analog. For this, we followed the same approach previously described. Briefly, exponentially growing *T. cruzi* epimastigotes were incubated with 100 µM EdU (Thermo Fisher Scientific) for 18 h at 27 °C. For each sample, an equivalent was prepared that was not incubated with EdU. Each sample was collected by centrifugation (1000 *g* for 7 min), washed twice using PBS, and fixed/permeabilized with 70% cold methanol overnight at 4 °C. The samples were vortexed, centrifuged, washed with PBS and incubated with 100 µL of Click-iT EdU detection solution for 1 h protected from light. The samples were then washed twice with PBS and resuspended in 500 mL of PBS. The cells were both analyzed in suspension using the BD Accuri C6 Flow Cytometer (BD Biosciences) and added onto microscope slides to be analyzed under a fluorescence microscope (Nikon Eclipse 80i). When analyzed by microscopy, Vectashield containing DAPI (VectorLabs) was used as an anti-fade mounting solution and to stain nuclei and kinetoplast DNA.

### Metabolic viability assay using 3-(4,5-dimethylthiazol-2-yl)-2,5-diphenyltetrazolium bromide (MTT)

MTT is a yellow tetrazole solution that is reduced to a purple formazan compound in metabolically active cells. Briefly, *T. cruzi* epimastigote forms from control and *IP6K*^-/+^ cells were started at a concentration of 2×10^6^ cells.mL^-1^ each. In the second day (exponential phase), the same volume (1 mL) for each sample was collected and placed into a 48 wells flat bottom plate. Next, the plate was centrifuged (1000 *g* for 7 min) and the medium was replaced with 3 mg.mL^-1^ MTT solution (diluted in PBS). Each sample containing parasites was resuspended in the MTT solution and incubated for 4 h with gentle shaking. Then, the samples were centrifuged (1000 *g* for 7 min), the MTT solution was discarded and the remaining formazan crystals were resuspended with 100% dimethyl sulfoxide (DMSO) (Sigma-Aldrich). The absorbance for each well plate was measured at 570 nm using a microplate spectrophotometer (Epoch – Biotek).

### Induction of metacyclogenesis

Metacyclogenesis was induced in Grace medium supplemented with 10% FBS. Briefly, *T. cruzi* epimastigote forms in exponential phase (2×10^7^ cells.mL^-1^) were collected by centrifugation (1000 *g* for 7 min) and washed twice with Grace medium. Next, the epimastigotes were settled in bottles containing 10 mL of Grace medium (initial 10^6^.mL^-1^) and kept for 10 days at 26 °C (Martínez-Díaz et al., 2001).

To verify the efficiency of metacyclogenesis, aliquots from the supernatant were collected by centrifugation on 5^th^ and 10^th^ days, washed twice with PBS, fixed in 4% paraformaldehyde (PFA) and settled in glass slides for immunofluorescence. Next, we used a mouse α-flagellin antibody (diluted 1:10 in 4% BSA solution) to recognize the *T. cruzi* flagellum (da Silva et al., 2013). The procedures for performing immunofluorescence were the same as those described previously. As a secondary antibody, we used Alexa Fluor 488 anti-mouse IgG (diluted 1:1000 in 4% BSA solution). 5 μL of VectaShield containing DAPI was added onto the slides, and a coverslip was mounted. Images were acquired using the fluorescent microscope Zeiss Axiovert 200M (Zeiss). It is worth highlighting that the position of the flagellum and organelles containing DNA (kinetoplast and nucleus) allows us to identify and estimate the percentage of *T. cruzi* cells in each stage of differentiation during metacyclogenesis (Bayer-Santos et al., 2013).

To double-check metacyclogenesis efficiency, we performed RT-qPCR to verify the expression of Metacyclin III (TcCLB.510943.44), a gene differently expressed in epimastigotes and trypomastigotes during metacyclogenesis. For this, the total RNA was extracted from 1x10^7^ parasites collected in 5^th^ and 10^th^ days of metacyclogenesis process according to the manufacturer’s instructions of Monarch^®^ Total RNA Miniprep Kit (New England Biolabs). Then, cDNA was obtained using SuperScript III (Invitrogen). Briefly, 0.8 ng/sample of RNA was used in the first reaction (8 µL RNA, 1 µL 50uM oligo-dT, 1 µL 10mM dNTPs) and incubated in 65 °C for 5 min, followed by ice 1 min. The second reaction was performed according to the manufacturer’s instructions (2 µL Buffer 10x RT, 4 µL of 25 mM MgCl_2_, 2 µL of 0.1 M DTT, 1 µL of RNase Out and 1 µL of Superscript III) and incubated at 20 °C for 50 min, 85 °C for 5 min, followed by addition of 1 µL of RNase H incubated at 37 °C for 20 min. The qPCR reaction was performed using Power Sybr Green Master Mix (Applied Biosystem): 5 µL of Sybr Green mix 2x, 1 µL of 2 µM of each Metacyclin III primer (forward and reverse) (see Supplementary Table 2), 20 ng cDNA and cycled in Step One Plus Real Time PCR System (Applied Biosystems).

### Infection assays using human cardiomyocytes AC16

For infection assays, AC16 human cardiomyocytes (5×10^4^ cells/well) were plated on circular glass coverslips placed in a 24-well plate and incubated overnight (∼16 h) for adhesion. Then, cells were infected with the TCTs forms from control and *IP6K*^-/+^ cells (MOI 10:1) and incubated at 37 °C in the presence of 5% CO_2_ for 3 h. After invasion, cells were washed to remove non-internalized parasites and fixed (using 4% PFA) for 15 min at 4 °C, in different time-points: 0 h – immediately post-washing (invasion analysis) – 24, 48, 72, 96, and 120 h post-invasion. The fixed samples were then washed with PBS and incubated overnight with human chagasic serum (diluted 1:1000 in blocking/permeabilizing PGN solution: 0.25% gelatin, 0.1% azide, and 0.5% saponin in PBS) at 4 °C. This serum was provided as a gift by prof. Renato Mortara (Federal University of São Paulo, Brazil). Next, the coverslips containing the cells were washed with PBS and incubated for 1 h with the following solution: secondary antibody Alexa Fluor 488 goat anti-human IgG (Thermo Fisher Scientific) (diluted 1:1000 in PGN solution), phalloidin-rhodamine (Invitrogen) (diluted 1:1000 in PGN solution), and 5 µg.mL^-1^ DAPI. After the incubation period (1 h), the coverslips containing the cells were washed twice with PBS and mounted in microscope slides using glycerol buffered with 0.1 M Tris, pH 8.6, with 0.1% p-phenylenediamine (PPD) (Sigma-Aldrich) as an anti-fade agent (Platt and Michael, 1983).

To determine the invasion and replication rates, the number of infected cells (total of 100 cells/replicate) and the number of amastigotes per cell (in the total of 100 infected cells/replicate) were analyzed, respectively. The images were obtained in a Confocal Microscopy (Leica SP8), using the software LASX office (Leica).

### Replication and quiescence analysis of egressed T. cruzi forms

The replication rate and quiescence of egressed parasites were analyzed using EdU as described before, with slight modifications. At 120 h post-infection, egressed parasites were collected and centrifuged (3000 *g* for 5 min), resuspended in 1 mL of DMEM medium and incubated with 10 µM of EdU for 1 h (DNA replication analysis) or 18 h (quiescence analysis). After incubation, parasites were fixed with 4% PFA for 15 min, washed with PBS and added into a 0.1% poli-L-lysine pre-treated slide (Knittel Glaser). The slides were then permeabilized with 0.5% Triton X-100 (Sigma-Aldrich) (diluted in PBS) for 5 min and briefly washed with PBS. Next, the parasites were labelled using the ‘click’ chemistry reaction and Vectashield containing DAPI (VectorLabs) was used as an anti-fade mounting solution and to stain nuclei and kinetoplast DNA, as previously described. The slides were analyzed by fluorescence microscope (Zeiss Axiovert 200M). For replication and quiescence analysis, 100 amastigotes/replicate were analyzed.

### Trypomastigogenesis

Cardiomyocytes (1×10^5^ cells/well) were plated on 6-well plate and incubated with TCTs (MOI 10:1) for 3 h. Next, cells were washed with PBS to eliminate non-internalized parasites and 96 h post-invasion, the cells were treated with 10 µM EdU for 24h. At 120 h post-invasion, the cells were detached using 300 µL/ well of trypsin-EDTA solution (T1757/ Vitrocel) and resuspended in 1mL of DMEM medium. The cells were physically ruptured using a 21-gauge needle coupled to syringe (5 mL), and intracellular parasites were recovered by differential centrifugation. Briefly, the total content was centrifuged at 200 *g* for 5 min. The supernatant was collected, placed into a new tube and centrifuged again at 3000 *g* for 5 min to recover the parasites. The parasites were fixed in 4% PFA (for 15 min at 4 °C), washed with PBS, and added into a 0.1% poli-L-lysine pre-treated slide (Knittel Glaser). The slides were then permeabilized with 0.5% Triton X-100 (Sigma-Aldrich) (diluted in PBS) for 5 min and briefly washed with PBS. Next, the parasites were labelled using the ‘*click’* chemistry reaction, as previously described. Finally, the cells were washed twice with PBS and 15 μL of glycerol buffered with 0.1 M Tris, pH 8.6, with 0.1% p-phenylenediamine (PPD) (Sigma-Aldrich) containing 5 µg.mL^-1^ DAPI, added onto the slides and a coverslip was mounted. The slides were fixed with nail polish before being observed under a fluorescence microscope (Zeiss Axiovert 200M). Different stages of trypomastigogenesis were defined according to the position of the nucleus, kinetoplast and flagellum. The parasites were classified into amastigotes, epimastigotes-like, intermediate forms and trypomastigotes (Tomasina et al., 2024). For trypomastigogenesis analysis, 200 cells per replicate were analyzed.

### RNA sequencing and transcriptome analysis

RNAs from each sample analyzed were isolated using the RNeasy Mini Kit (Qiagen) and treated with RNAse-free DNAse I (Qiagen). RNA samples were quantified and evaluated for integrity using the fluorometer Qubit 4 (Thermo Scientific). All samples showed an A260/A280 ratio close do 2.0.

Standard libraries for massive sequencing were generated using the TruSeq RNA Sample Prep Kit (Illumina). Briefly, poly-A + RNA was selected by oligo-dT chromatography, and RNA fragmentation was achieved using divalent cations under elevated temperature. Afterwards, these fragments were used to generate a cDNA library, and cDNA fragments corresponding in size to about 450–550 bp were isolated from an agarose gel. Single-indexed primers were added in a Combinatorial Dual (CD) index process. Paired-end reads of 150 bp (30 M reads per sample) were obtained from Beijing Genomics Institute (BGI), Shenzhen, China.

NGS raw data (.fastq files) were preprocessed using the software fastp (version 0.20.1). The trimmed files were then mapped to the CL Brener Non-Esmeraldo like genome sequence (release 68), obtained from the TriTrypDB database (Aslett et al., 2009) using the bowtie2 software. To assess the separation between replicates from each group (Epimastigotes and Trypomastigotes – TCTs), principal component analysis (PCA) was performed using the prcomp function (Supplementary Figure 4A). Differential gene expression between groups was conducted using edgeR (Robinson et al., 2009) (version 3.36.0). The samples from the groups Control (trypomastigotes – TCTs) vs. *IP6K*^-/+^ (trypomastigotes – TCTs) and control (epimastigotes) vs. *IP6K*^-/+^ (epimastigotes) were compared. Before the comparison, both count matrices were filtered to only include expressed genes, using the filterByExpr function from the edgeR package. The average percentage of mapped reads was 73% and the quantified reads was (Supplementary Figure 4B, C). The .bam files were quantified using featureCounts software (version 2.0.3) (Liao et al., 2014) to generate a count matrix containing 11,106 detected genes, corresponding to 57% of the genes annotated in the CL Brener Non-Esmeraldo like genome.

It is worth mentioning that since the number of TCTs released 120 h after infection was very low in the *IP6K*^-/+^ lineage, we had to perform the trypomastigogenesis process several times until we obtained a significant amount of TCTs to perform RNA extraction. We chose to group these recovered TCTs into a single pool and therefore we had only a single sample of TCTs for transcriptomics analysis (Supplementary Figure 4A). We recognize that using a single replicate limits the statistical power for this condition. However, we chose to prioritize data quality over including a lower-quality dataset due to the small sample size, which could introduce bias or noise into the analysis. Due to this, an arbitrary value was assigned for the dispersion estimate, with a coefficient of variation (BCV) of 0.1 fixed for calculating dispersion between samples of this group, as recommended by the edgeR manual. Differential expressions were analyzed using the exact test, and genes with an adjusted p-value (FDR) ≤ 0.05 and logFC |0.58| were considered differentially expressed. For the epimastigotes group, we used FDR ≤ 0.05 and logFC |0.58| as criteria.

Of note, the distance between the epimastigote replicates in the PCA plot may, at first glance, indicate variability. However, this probably reflects technical variation associated with independent RNA-seq library preparations derived from separate cell culture batches (biological replicates), rather than fundamental biological divergence. To further assess these data robustness and confirm this, we performed pairwise correlation analyses across all samples and verified that correlations between biological replicates were consistently higher than those observed between control and KOs samples, or between Epimastigotes and TCT samples (Supplementary Figure 4D), supporting the overall consistency of the dataset and its suitability for downstream comparative analyses.

Functional enrichment analysis was conducted using topGO (Alexa and Rahnenfuhrer, 2024) and gene ontologies from Cellular Components, Biological Processes and Molecular Function retrieved from TriTrypDB database. Terms with enrichment displaying p ≤ 0.05 as determined by a Kolmogorov-Smirnov test were considered as significant. A similar analysis was performed using the enrichment tool hosted by TriTrypDB. For this, the list of differentially expressed genes in each group was used as input, and terms that were computed or curated and had a Bonferroni-corrected p-value ≤ 0.05 were considered enriched.

### Data availability

The WGS raw data have been deposited in the NCBI Sequence Read Archive (SRA) under BioProject PRJNA1332607 (accessions: SAMN51753950, SAMN51753951 and SAMN51753952 for *T. cruzi* WT, T7/Cas9 and *IP6K*^-/+^, respectively). The raw and processed RNAseq data have been deposited in the Gene Expression Omnibus (GEO) under accession GSE309901.

### Statistics and reproducibility

GraphPad Prism 9.0.0 package (GraphPad Software, Inc.) was used for the most statistical analysis carried out in this work. All datasets were checked for normal distribution using a Shapiro–Wilk normality test. Significant differences between groups were identified using two-tailed Student’s t-test (paired or unpaired, as appropriate). Comparisons of more than two groups were performed with one-way ANOVA followed by Student–Newman–Keuls test. Data is represented as mean ± SEM (standard error of the mean), unless otherwise stated.

## Results

### IP6K is slightly conserved among eukaryotes and presents low sequence similarity relative to its human homologs

To investigate IP6K conservation and similarity among eukaryotes, we carried IP6K phylogenetic analysis encompassing the most important eukaryotic groups (Figure 1A). Phylogenic trees were inferred using RAxML-NG’s Maximum Likelihood, whereby 706 amino acid sequences from IP6K of different eukaryotes (one representative isoform for each organism) were analyzed. All IP6Ks were examined for the presence of the PxxxDxKxG motif (Supplementary Figure 1A), which is a conserved signature across the inositol phosphate kinases (IPK) family (Wang et al., 2014).

**Figure 1.**
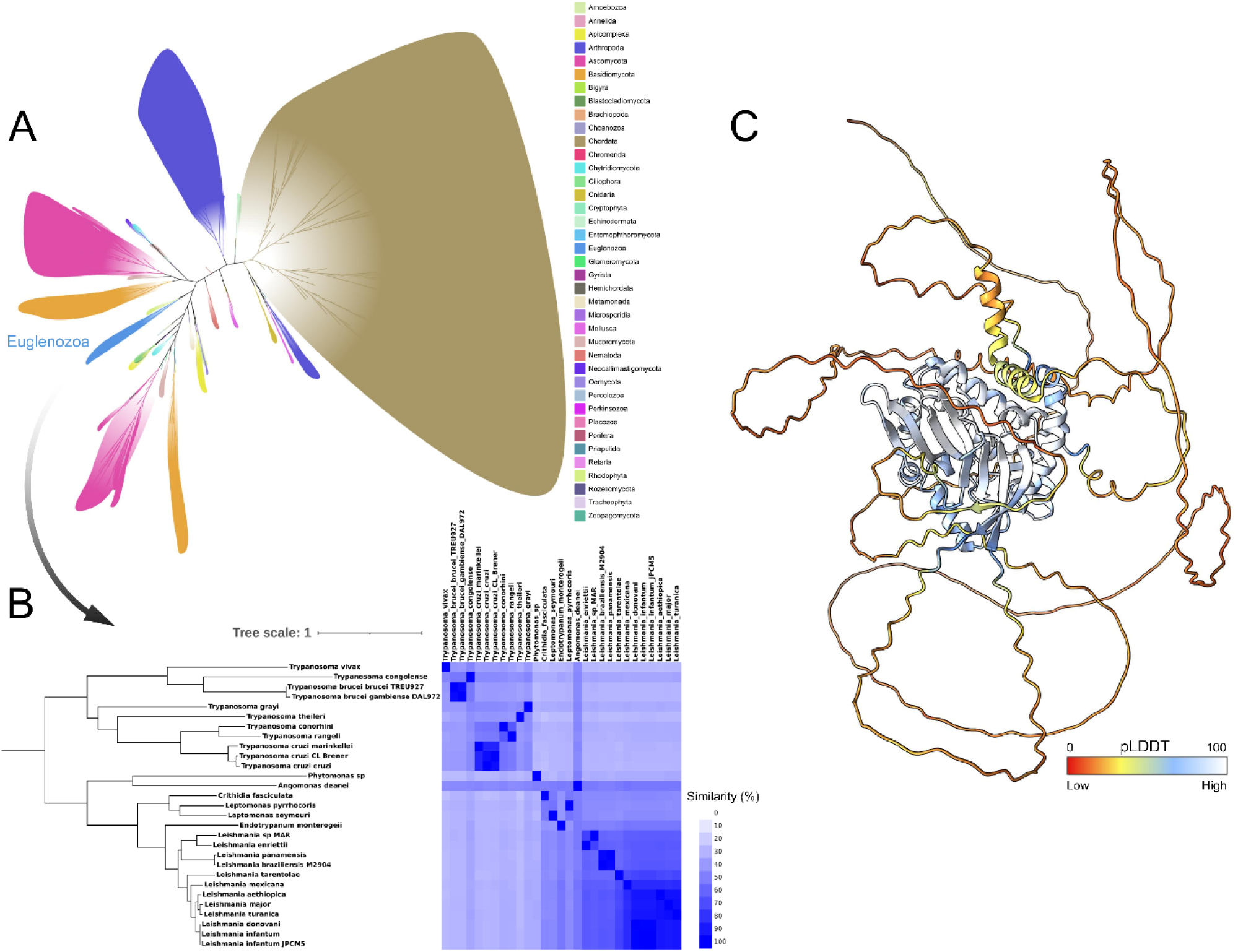
*In silico* analyses of IP6K suggest slight conservation of this kinase among non-plant eukaryotes. **A**. IP6K phylogenetic analysis of 706 representative IP6K orthologs constructed on Maximum Likelihood with “JTT+G” matrix rate and 650 bootstraps. Organisms from the Euglenozoa family are highlighted in light blue. **B.** IP6K sequence similarity heatmap mapped showing comparisons of members from Euglenozoa. **C**. Tertiary structure of *T. cruzi* IP6K, as predicted by AlphaFold3. pLDDT (predicted local distance difference test) is a per-residue measurement of local confidence. Higher scores indicate higher confidence (accurate prediction).

According to our results, the classification of the different IP6K orthologs into different clades suggests a divergent evolutionary path, which may have contributed to the emergence of moonlighting functions for this kinase (Figure 1A). Our analysis confirms, as expected, that IP6K is absent in most plant species (Figure 1A), since the inositol tetrakisphosphate kinase (ITPK) family can fulfill the IP6K role (IP_7_ synthesis) in this group (reviewed in Mihiret et al., 2024). These data corroborate with other studies and suggest that IP6K is not essential for IP_7_ synthesis, having redundant functions with other kinases (reviewed in Williams et al., 2015; Mihiret et al., 2024).

Within Euglenozoa, the phylogenetic group that encompass trypanosomatids (Adl et al., 2012), IP6K retained moderate similarity among trypanosomatids: *T. cruzi* CL Brener strain shows 99.73% similarity (E-value of 0.0) with *T. cruzi* Y strain, 56.27% similarity (E-value of 2^-82^) with *Trypanosoma brucei* (TREU927), and 47.62% similarity (E-value of 2^-60^) with *Leishmania* (*L*.) *donovani* (Figure 1B).

Additionally, our findings suggest low conservation between *T. cruzi* IP6K and distantly related orthologs from other eukaryotes, especially the human isoforms. *T. cruzi* IP6K shows 26.92% similarity (E-value of 3^-18^) with human IP6K isoform 1 (*Hs*_IP6K1), 25.21% similarity (E-value of 2^-17^) with human IP6K isoform 2 (*Hs*_IP6K2), and 25.07% similarity (E-value of 2^-21^) with human IP6K isoform 3 (*Hs*_IP6K3) (Supplementary Figure 1A). In general, these low levels of conservation between *T. cruzi* IP6K and its human homologs represent a favorable feature for drug development efforts (Schmidt et al., 2021; Baker et al., 2021).

Although the PxxxDxKxG motif is conserved, *T. cruzi* IP6K also possesses a large unique N-terminal region (residues 1–299) that could confer specific or exclusive functions (Supplementary Figure 1A). The predicted tertiary structure of *T. cruzi* IP6K shows the N-terminal portion to be largely disordered, although this may in part reflect modeling uncertainty (Figure 1C). Nevertheless, it is possible to infer that the highly conserved PxxxDxKxG motif and some other amino acid residues (Arg626, His340, Lys588, Lys591, and Ser 587) are apparently essential for the IP_7_ synthesis from IP_6_ and ATP, in the presence of 2 Mg^2+^ ions (Supplementary Figure 1B).

### The loss of a single IP6K allele impair proliferation but does not significantly affect inositol pyrophosphate levels in T. cruzi

To investigate functions of *T. cruzi* IP6K, we performed a loss-of-function approach using the CRISPR/Cas9 system in epimastigote forms (see material and methods). The first round of transfections was performed using a donor DNA containing a blasticidin-resistance gene. PCR results of the nine clones selected confirmed the presence of the blasticidin-resistance gene inserted at the correct locus. However, our results also showed the other copy of *IP6K* in its correct locus, evidencing that all clones selected were single-allele mutants for *IP6K* (IP6K^-/+^) (Supplementary Figure 2B). The inability to perform a complete knockout of this gene in one round of transfection seems to reflect on the hypothesis that *IP6K* is an essential gene for *T. cruzi* (Mantilla et al., 2021). However, we persisted in attempting to disrupt the remaining *IP6K* copy using a donor DNA containing a puromycin-resistance gene and transfecting one of the previously selected single-allele *IP6K* mutant clones (clone 2) from the first round of transfection.

PCR analysis (2^nd^ day of the growth curve) revealed both puromycin- and blasticidin-resistance cassetes correctly integrated at the *IP6K* target loci, consistent with the generation of a *IP6K* null mutant (Supplementary Figure 2C, primers F1+R3 and F1+R4). However, amplification of *IP6K* allele was also detected (Supplementary Figure 2C, primers F2+R2 and F1+R2). This likely occurred because *IP6K* null mutants (*IP6K*^-/-^) were unable to proliferate, and the cell culture at that time represented a mixed polyclonal population (Supplementary Figure 2D-E). In this context, genomic DNA from single-allele mutants (*IP6K*^-/+^) could still be present in the culture, masking the results. The inability of parasites lacking *IP6K* to survive not only explains the absence of a *IP6K^-/-^*lineage after the first round of transfection but also further supports *IP6K* as an essential gene in *T. cruzi* (Mantilla et al., 2021). Future studies using conditional knockout systems, such as riboswitch-or DiCre-based approaches, could further challenge this hypothesis (Duncan et al., 2016; Lander et al., 2020; Yagoubat et al., 2020).

We then moved forward using IP6K^-/+^ cells to investigate the roles of this kinase in *T. cruzi*. We first verified the proliferation capacity of *IP6K*^-/+^ epimastigotes from clone 2 relative to parental T7/Cas9 (Control) and wild-type (WT) *T. cruzi*. As expected, the WT (green) and parental T7/Cas9 – Control (gray) groups did not show significant differences between them (Figure 2A). However, IP6K^-/+^ cells (blue) showed a slight but significant (*p* < 0.001) growth delay relative to the controls (T7/Cas9 and WT).

**Figure 2.**
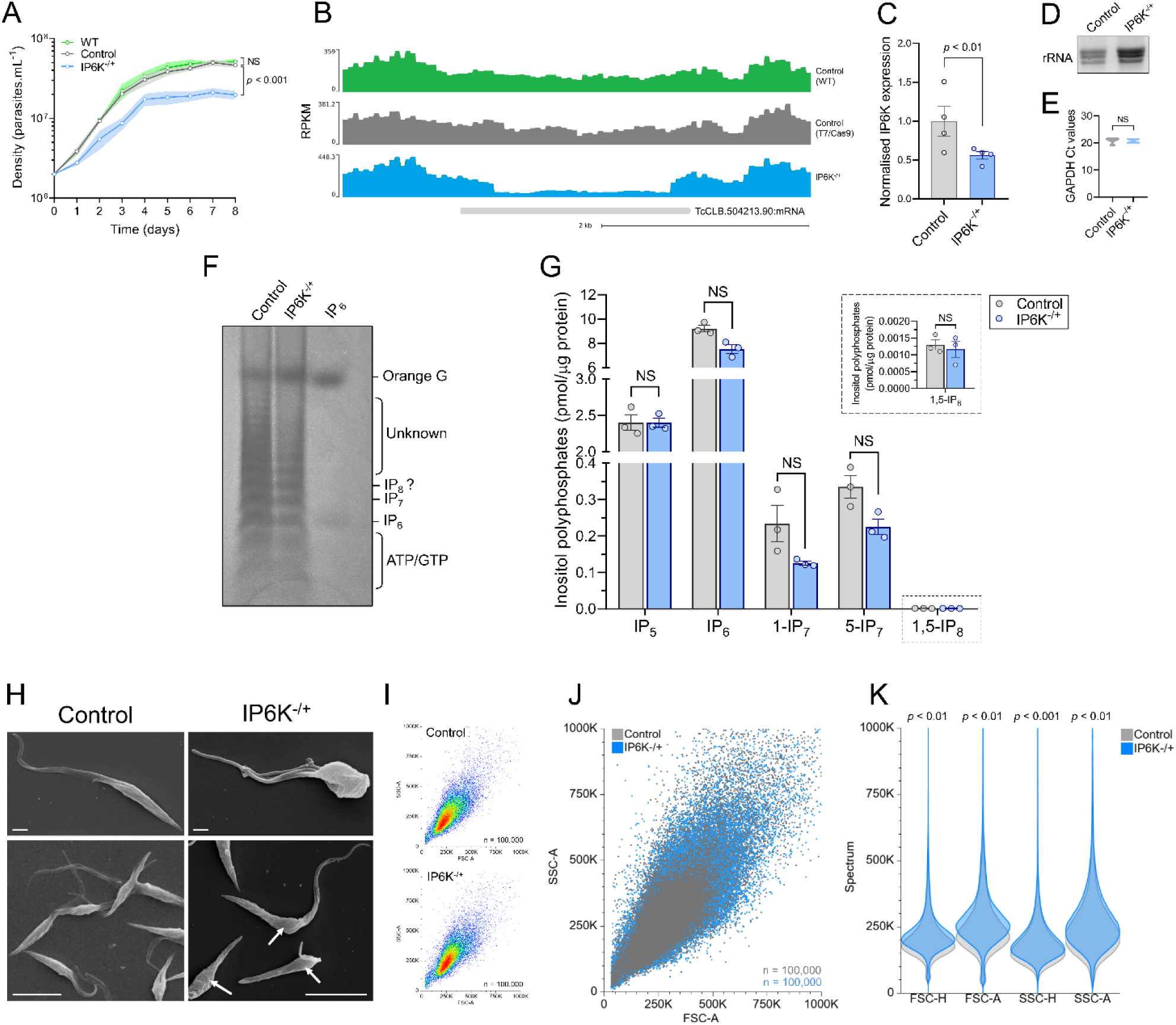
The single *IP6K* allele depletion impairs proliferation and alters *T. cruzi* epimastigote cell morphology. **A.** Growth curves showing proliferation patterns of WT, T7/Cas9, and *IP6K*^-/+^ cells. The shaded area represents the standard deviation (SD) of the mean values. **B**. A density plot of RPKM (reads per kilobase per million mapped reads) for *IP6K* locus for the WT (green), T7/Cas9 (gray), and *IP6K*^-/+^ (blue) cells. **C**. RT-qPCR results showing the normalized expression of *IP6K* in the control (T7/Cas9) and *IP6K*^-/+^ cells. The error bars represent the standard error of the mean (± SEM) from an assay performed in triplicate. **D**. Total RNA (1 μg) pattern used in RT-qPCR assays. The three bands evidenced in 1.2% agarose gel represent the different subunits of ribosomal RNA (rRNA). **E**. Ct values for GAPDH and *IP6K*^-/+^, obtained from quadruplicated assays. **F.** PAGE analysis of inositol polyphosphates extracted from *T. cruzi* epimastigote forms. IP6 (also known as phytic acid or inositol hexakisphosphate) was used as a molecular standard. **G.** CE–ESI–MS analysis of inositol polyphosphates levels (IP5, IP6, 1-IP7, 5-IP7, and 1,5-IP8) in Control and IP6K^-/+^ cells, normalized by the total amount of protein. The error bars represent the standard error of the mean (± SEM) from an assay performed in triplicate. NS = non-significant according to Student’s *t*-test. **H**. Scanning electron microscopy (SEM) analysis shows morphological alterations in the *IP6K*^-/+^ cells. **I**. Density plot of the size (FSC-A) and internal complexity (SSC-A) **J**. Merge of the density plots shown in *G*. **K**. Light spectrum measured for each parameter analyzed: FSC-height, FSC-area, SSC-height, and SSC-area.

To confirm the loss of a single *IP6K* allele and to mitigate the possibility that the CRISPR/Cas9 approach used might have generated unintended DNA breaks (off targets) that could contribute to the observed phenotype, we performed the whole genome sequencing (WGS) of the controls (T7/Cas9 and WT) and *IP6K*^-/+^ cells. Figure 2B shows the RPKM (reads per kilobase per million mapped reads) of the *IP6K* locus for the WT (green), T7/Cas9 (gray) and *IP6K*^-/+^ (blue) cells. The decrease in the number of reads only for the *IP6K*^-/+^ cells (Figure 2B) confirms that the expected, precise genome editing (single *IP6K* allele depletion) was successfully achieved.

To check for the presence of off-targets in the *IP6K*^-/+^ cells, we looked for deletions at two scales: a coarse whole-genome view (20 kb windows) and a finer pass (1 kb windows). In the coarse view, *IP6K*^-/+^ showed no clear sequence losses when compared with WT after correcting for overall depth differences. A few small read depth ‘dips’ on chromosome 31 were seen, but these occurred in repeat-rich regions where short reads do not map uniquely and were not supported by breakpoint patterns. Of note, they also overlapped changes presented in the parental T7/Cas9 cells. After applying pre-set filters, no 20 kb deletions remained that were specific to *IP6K*^-/+^. We then repeated the search with 1 kb windows to capture smaller events. After removing regions shared with T7/Cas9 and filtering out assembly joins and low-mappability areas, none of the preliminary intervals met evidence thresholds. Together, our WGS data indicates no detectable off target deletions in *IP6K*^-/+^, either at large (tens of kilobases) or small (kilobase) scales (Supplementary Table 6). As the WT (green) and T7/Cas9 (gray) cells did not exhibit significant differences in growth curves or WGS, we opted to use the parental T7/Cas9 lineage as the more appropriate control for the onward assays.

To confirm decreased *IP6K* transcript levels in the *IP6K*^-/+^ cells, we carried out RT-qPCR using *GAPDH* as the endogenous control (Figure 2C). Figure 2D shows the quality of the total RNA used for the RT-qPCR assays and Figure 2E presents violin plots of the Ct values for *GAPDH* in the control and *IP6K*^-/+^ cells. As expected, the *IP6K* transcript levels significantly reduced (*p* < 0.01) to approximately half of the control level (Figure 2C).

However, although the IP6K transcript levels were reduced as expected, the PP-IP levels (1-IP_7_, 5-IP_7_ and 1,5-IP_8_) did not show a significant reduction (Figure 2F, G). This finding corroborates the data obtained by Mantilla et al., (2021) and indicates that only one IP6K allele is sufficient to maintain the necessary PP-IP levels in epimastigote forms of *T. cruzi*. Furthermore, the maintenance of PP-IP levels in IP6K-deficient cells suggests that any differential phenotypes may not be directly related to PP-IPs, but rather to IP6K itself. Of note, we chose to normalize the levels of PP-IPs obtained by CE–ESI–MS analysis by the amount of protein extracted from each sample (Figure 2 G) instead of by the number of cells (Supplementary Figure 2F).

### Reduced IP6K levels impair cell morphology of T. cruzi epimastigote forms

Interestingly, when we observed the *IP6K*^-/+^ cells under the microscope, we noted subtle morphological alterations, with an apparent accumulation of cells with a single flagellum and reduced mobility. To further investigate these alterations, we used scanning electron microscopy (SEM). In contrast to the typical epimastigote forms (control), the *IP6K*^-/+^ cells showed a rounding of the cell body, especially at the anterior region near the flagellum (arrows in Figure 2H – *IP6K*^-/+^). Furthermore, *IP6K*^-/+^ also exhibited a shortening and wrinkling of the cell body relative to the control (Figure 2H – right top panel). To quantify these morphological alterations, we performed flow cytometry analyses to evaluate the relative cell size and internal complexity. We observed significant alterations in forward scatter (FSC-A and FSC-H) and side scatter (SSC-A and SSC-H) parameters, which represent changes in size and internal complexity, respectively (Figures 2I, J). The electromagnetic spectrum scattered from each analyzed cell was measured for each parameter (Figure 2K), and the median of each replicate was used for statistical analyses. Together, these findings suggest that loss of a single *IP6K* allele induces morphological alterations that may be correlated with reduced mobility and, consequently, proliferation impairment of *T. cruzi* epimastigote forms.

### Reduced IP6K levels cause a discrete arrest in G0/G1 and slightly alter the cell cycle phase lengths

To begin investigating the possible presence of cell cycle alterations in the *IP6K*^-/+^ cells that could result from morphological alterations and contribute to the proliferation impairment previously observed, we measured FKN (flagellum, kinetoplast, and nucleus) patterns in individual cells. The FKN analysis is commonly used to assess alterations in cell morphology and cell cycle progression in trypanosomatids (Marin et al., 2018), as it allows inference of distinct cell cycle phases: G1/early S (1F1K1N), late S (2F1K1N), G2/mitosis (2F2K1N), and cytokinesis (2F2K2N). Furthermore, abnormal/aberrant cells can be identified by quantifying FKN patterns that do not fit these typical configurations.

Using this approach, we found that IP6K^-/+^ cells showed a higher number of cells displaying the 1F1K1N pattern relative to the control, suggesting a potential cell cycle impairment (Figure 3A, B). To investigate this further, we performed DNA content analyses by flow cytometry using PI. We observed that *IP6K*^-/+^ (blue) exhibited a mild increase in the percentage of cells arrested in G1/S relative to the control (gray) (Figure 3C).

**Figure 3.**
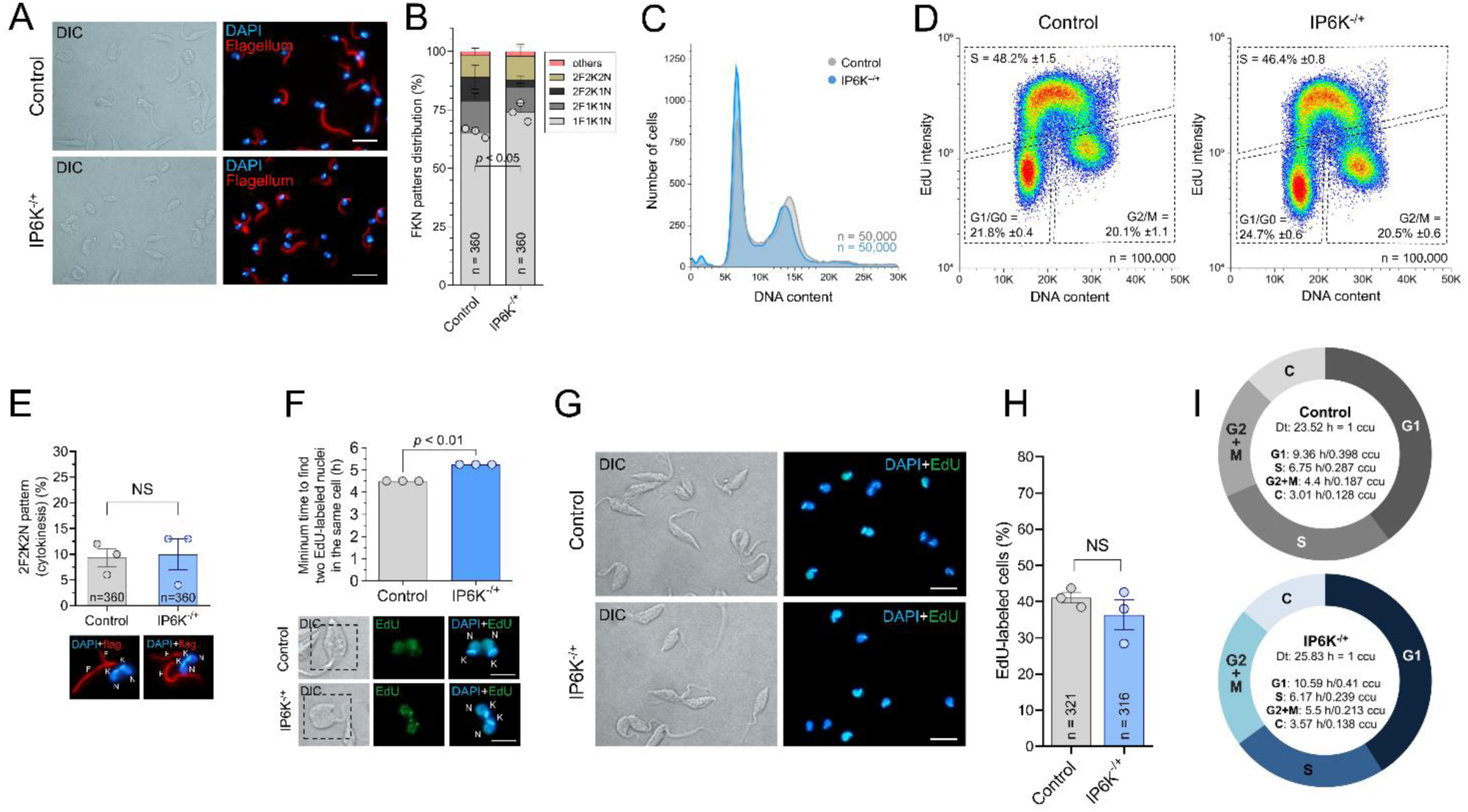
*IP6K*^-/+^ cells show a discrete arrest in G0/G1 and slight alterations in the cell cycle phase lengths. **A**. IFA analysis showing representative images evidencing the different FKN patterns found in control and *IP6K*^-/+^ cells. **B**. Quantification of the FKN patterns showed in A. The data represents the SD of three independent experiments. 360 cells were analyzed per group. **C**. Histograms overlay of the DNA content analysis using PI. **D**. Density plots of the double-labeled approach using DAPI and EdU. Hot colors (red and yellow) indicate areas of higher cell density, while cooler colors (green and blue) indicate areas of lower cell density. These assays were carried out in biological triplicate, and 100,000 events were analyzed per sample. **E**. Quantification of cells in the cytokinesis phase (2F2K2N) determined by the FKN patterns analysis made in A. **F**. (Top) Minimum time to find two EdU-labeled nuclei within the same cell. (Bottom) Representative images used to estimate the time in F (Top). **G**. Representative images of the EdU-labelled cells after 1 h EdU pulse. **H.** Quantification of the EdU-labeled cells shown in G. **I.** Estimation of the cell cycle phases duration performed using the CeCyD software (https://cecyd.vital.butantan.gov.br/) (da Silva et al., 2020). The doubling time for the two lineages were estimated by taking the values at exponential phase and using Doubling Time software (http://www.doubling-time.com).

To discriminate whether this arrest occurred in G0/G1 or early S phase, we performed a double-labeled approach using DAPI staining and a 30 min EdU-pulse to monitor DNA replication. We observed that *IP6K*^-/+^ cells showed a higher percentage of cells in G0/G1 (24.7% ±0.6) compared to the control (21.8% ±0.4) (Figure 3D), which could be generated either by DNA damage or increased G0 state (quiescent) cells.

If we assume that the main function of IP6K is IP_7_ synthesis and that PP-IPs levels fluctuate dynamically throughout the cell cycle phases, even without altering their total availability (Barker et al., 2004), the observed partial G0/G1 arrest may reflect PP-IPs reduced availability. In yeast, it has been suggested that PP-IPs synthesized by KCS1 (the *IP6K* homolog in Fungi) modulate S phase progression following a G0/G1 cell cycle arrest. However, it remains unclear whether, in addition to modulating S-phase progression, PP-IPs also contribute to G0/G1 arrest in these organisms (Banfic et al., 2013; Banfic et al., 2016). Thus, determining the duration of each cell cycle phase and assessing the S-phase progression will clarify whether the S-phase modulation observed in yeast is conserved in *T. cruzi*.

To estimate the duration of the cell cycle phases and simultaneously evaluate S-phase progression, we used a previously standardized approach (da Silva et al., 2020). First, we measured the following parameters: doubling time (estimated from the growth curve – Figure 2A), percentage of cells in cytokinesis (Figure 3E), minimum time required to observe two EdU-labeled nuclei within the same cell (Figure 3F), and percentage of EdU-positive cells after 1 h EdU pulse (Figure 3G, H) (da Silva et al., 2020). We then applied calculations based on these parameters to estimate the duration of each cell cycle phase duration in both *T. cruzi* groups (control and *IP6K*^-/+^) (Figure 3I). For this, we used the CeCyD software (https://cecyd.vital.butantan.gov.br/) (da Silva et al., 2020).

We found subtle changes in the cell cycle duration of *IP6K*^-/+^ relative to the control, including a slight increase in the G1 phase. This finding is consistent with the FKN pattern (Figures 3A, B) and DNA content analyses (Figures 3C, D), supporting the hypothesis that *IP6K* is required for proper progression through the cell cycle phases. However, although S-phase duration in IP6K^-/+^ cells was slightly shorter (Figure 3I), the S-phase progression itself was not significantly impaired (Figure 3D and H), in contrast to what has been reported in yeast (Banfic et al., 2013; Banfic et al., 2016).

### Reduced IP6K levels lead part of the T. cruzi epimastigotes population to quiescence

In general, two possibilities could account for the partial cell cycle arrest observed at G0/G1: the presence of DNA damage or an increase in the number of cells with undamaged DNA but non-committed with the cell cycle (i.e., quiescent cells). To investigate these possibilities, we performed assays designed to discriminate between these two scenarios.

The cellular decision to progress through the cell cycle is mediated by a specific regulatory machinery that responds to various internal and external signals (Pack et al., 2019). Moreover, distinct mechanisms act in concert to trigger cell cycle arrest, usually in response to DNA damage (Toettcher et al, 2009). Therefore, the most parsimonious explanation for a partial cell cycle arrest in the *IP6K*^-/+^ cells is the presence of DNA damage. To test this hypothesis, we carried out immunofluorescence assays (IFA) using an α-γH2A antibody (Figures 4 A, C) and TUNEL analysis (Figures 4 B, D) to investigate the presence of DNA damage in the analyzed lineages. Phleomycin was used as a positive control for γH2A detection (Sleigh, 1976; Glover and Horn, 2012), and DNase I served as a positive control for TUNEL assay (da Silva et al., 2017) (Figures 4 A-D).

**Figure 4.**
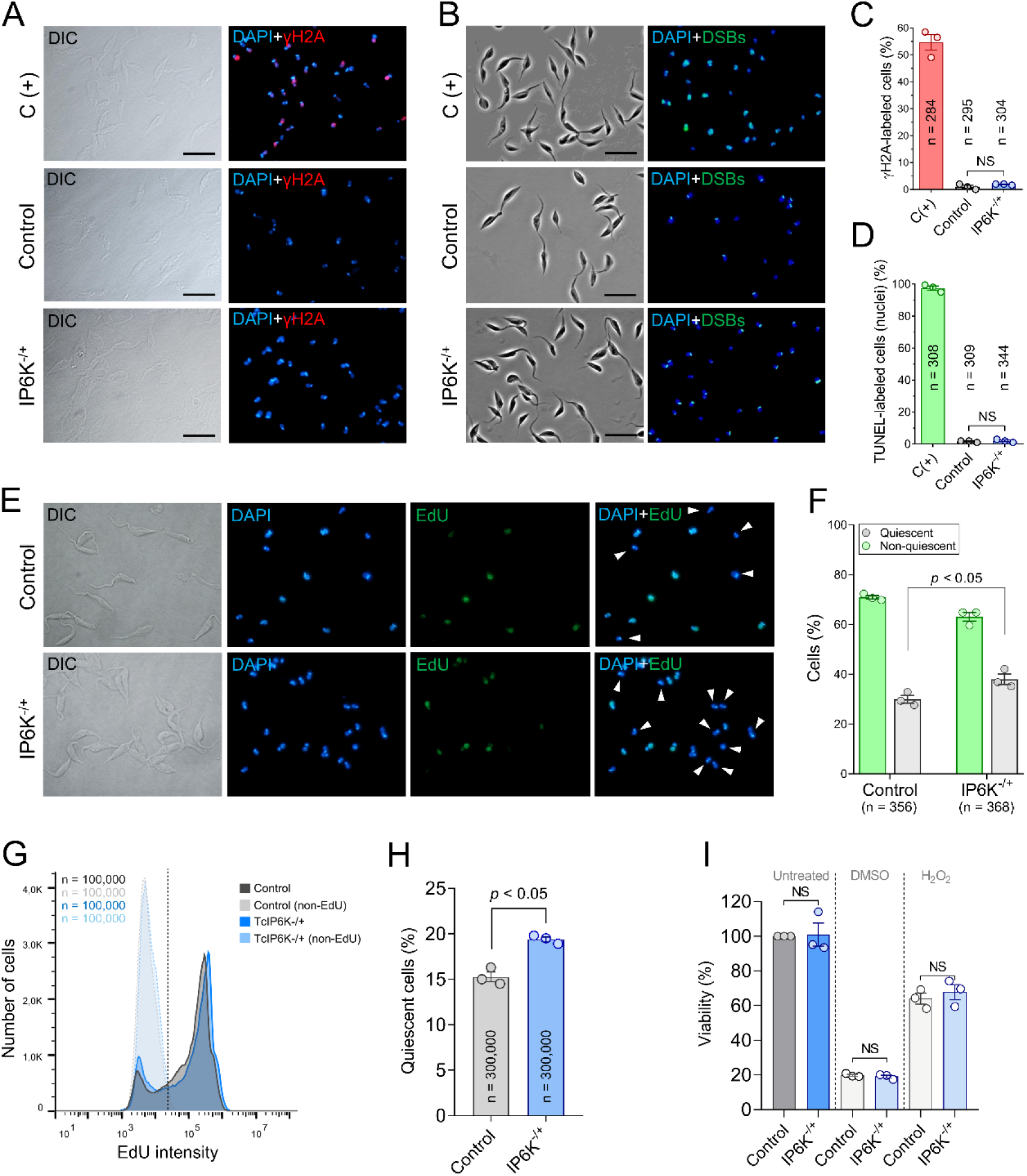
The loss of a single *IP6K* allele leads *T. cruzi* epimastigotes to quiescence. **A**. Representative images of IFA evidencing the absence of γH2A foci in the control and *IP6K*^-/+^ cells. The positive control was carried out pre-treating the control with phleomycin. Bars = 5 μm. **B**. Representative images of the TUNEL assay evidencing the absence of DNA breaks (DSBs) in the control and *IP6K*^-/+^ cells. The positive control was carried out pre-treating the control sample with DNase I. **C**. Quantification of the cells presenting γH2A foci. **D**. Quantification of the cells presenting DSBs foci. The error bars represent the standard error of the mean (± SEM) from an assay performed in triplicate. NS = non-significant according to Student’s *t*-test. **E.** DNA replication monitoring (after 18 h EdU) evidencing that *IP6K*^-/+^ cells presents more non-labeled cells (quiescent) (white arrows) relative to the control. **F**. Quantification of the EdU-labeled and non-labeled cells. The error bars represent the standard error of the mean (± SEM). **G**. Overlapped histograms of the control (gray) and *IP6K*^-/+^ (blue) cells. The same lineages were cultivated in the absence of EdU, being used as non-labeled control [control (light gray) and *IP6K*^-/+^ (light blue)]. 100,000 cells were analyzed per sample in an assay performed in triplicate. **H**. Measurement of the EdU-labeled and non-labeled (quiescent) cells presented in G. The error bars represent the standard error of the mean (± SEM) from an assay performed in triplicate. **I**. Viability assay shows that the difference in EdU uptake observed was not due to an imbalance in viability.

Surprisingly, neither technique revealed detectable DNA damage in the analyzed lineages (Figures 4 A, D). This finding prompted us to test the second hypothesis to explain the G0/G1 arrest observed in our results: an increased number of quiescent cells. Quiescence is a state of reversible proliferative arrest in which cells are not committed to division but still retain the capacity to reenter the cell cycle upon receiving an appropriate stimulus (Marescal and Cheeseman, 2020). To quantify quiescent cells, we used an approach based on EdU-negative selection, considering the previously estimated cell cycle phase duration (Figure 3I).

Our findings allowed us to identify fewer EdU-labeled cells in *IP6K*^-/+^ (60.5 ±1.99) relative to the control (67.3 ±2.99), consequently, more quiescent (EdU non-labeled) cells (38.44 ±4.8 in *IP6K*^-/+^ relative to 30.26 ±6.7 in the control group) (Figures 4E, F). By flow cytometry, the same lineage presented a slightly higher non EdU-labeled peak (blue peak on the left side), consistent with the IFA (Figures 4G, H). A possible explanation for these data is that EdU-negative cells are not viable, rather than being non-replicating. To test this possibility, we performed a viability assay (Figure 4I). The efficiency of this assay was demonstrated using two positive controls: Dimethylsulfoxide (DMSO) and hydrogen peroxide (H_2_O_2_), which cause most of the population to die, decreasing the viability of both groups (Figure 4I). When we observed the non-treated groups, our results showed that *IP6K*^-/+^ cells had the same viability as the control group (Figure 4I), indicating that the EdU-negative cells observed previously (Figures E-H) are living, suggesting an increase in quiescent cells following the loss of a single *IP6K* allele.

### Reduced IP6K levels impair metacyclogenesis

*T. cruzi* metacyclic trypomastigotes differentiate (*in vitro*) normally between the fifth and tenth day of a standard epimastigote growth curve, and are naturally quiescent (Christiano et al., 2017; Martín-Escolano et al., 2022). Thus, a possible explanation for the increased quiescent population in the *IP6K*^-/+^ cells could be the presence of early-differentiating metacyclic trypomastigotes.

To verify the presence of metacyclics, we stimulated metacyclogenesis by culturing control and *IP6K*^-/+^ epimastigotes in Grace’s medium for 10 days (initial 10^6^.mL^-1^). The efficiency of metacyclogenesis was checked by collecting and analyzing aliquots on the 5^th^ and 10^th^ days (Figure 5). Through the location of the kinetoplast relative to the nucleus, kinetoplast shape and kDNA topology (Figure 5A), it is possible to easily distinguish epimastigotes, intermediates and metacyclic forms during the metacyclogenesis (Gonçalves et al., 2018). Of note, *T. cruzi* intermediate forms during metacyclogenesis are characterized by progressive elongation, posterior kinetoplast movement, and nuclear changes that lead to the infective metacyclic trypomastigote form (Gonçalves et al., 2018).

**Figure 5.**
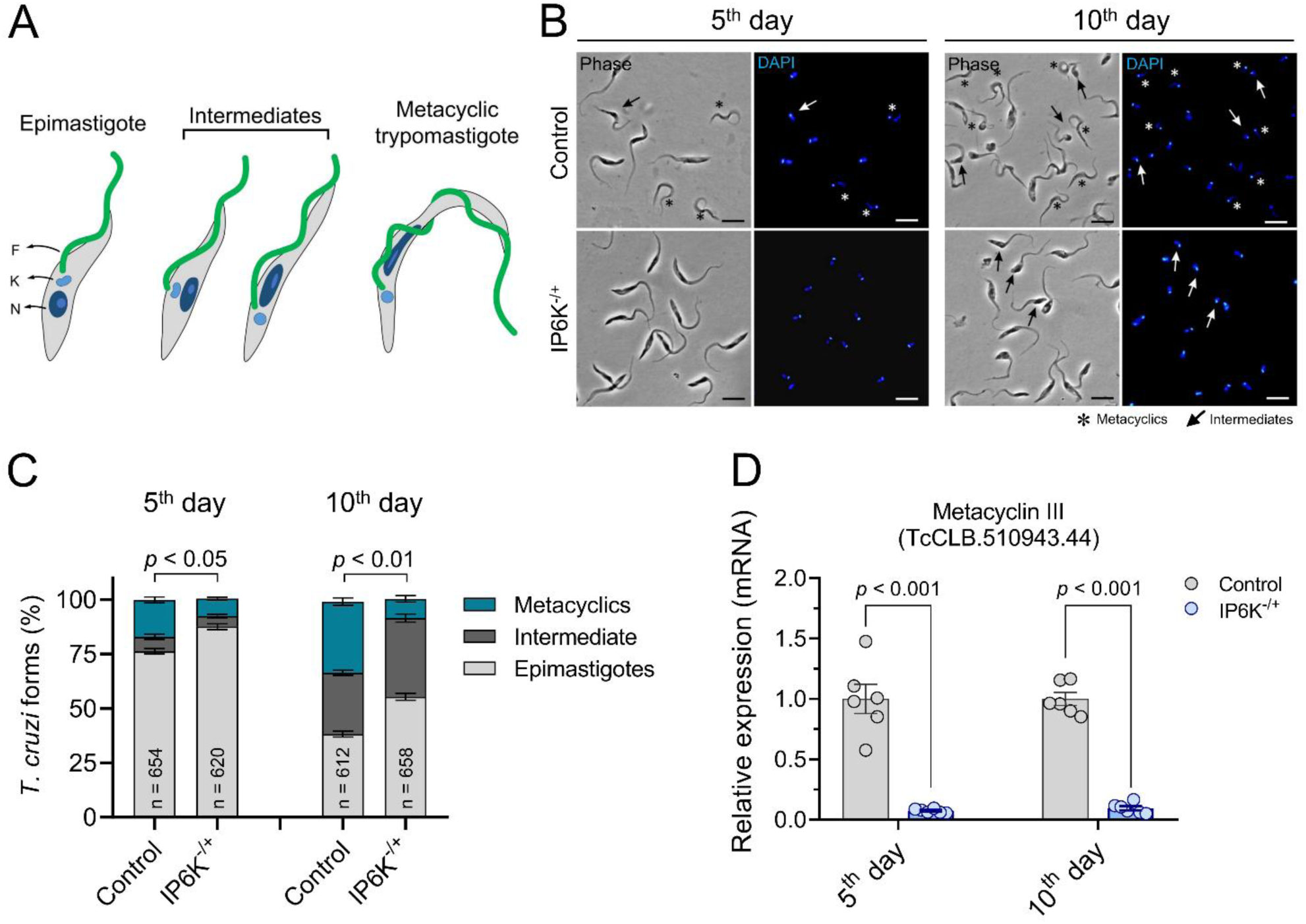
The loss of a single *IP6K* allele impairs metacyclogenesis. **A**. Schematic representation of *T. cruzi* life forms during metacyclogenesis. **B.** Representative images of parasites in different stages of metacyclogenesis. Phase contrast (gray) was used to depict cell morphology. DAPI (blue) was used to stain DNA-containing organelles (nucleus and kinetoplast). Arrows indicate intermediate forms, asterisk indicates metacyclic forms. Bars = 5 µm. **C**. Percentage of parasites at each differentiation stage after the fifth and tenth days of culturing in Grace’s medium. The error bars represent the standard error of the mean (± SEM) from an assay performed in triplicate. **D**. RT-qPCR results showing the normalized expression of Metacyclin III (TcCLB.510943.44) in the control (T7/Cas9) and *IP6K*^-/+^ cells. The error bars represent the standard error of the mean (± SEM) from an assay performed in sextuplicate.

Our finding revealed that *IP6K*^-/+^ cells showed less metacyclic forms relative to control during metacyclogenesis kinetics – 5^th^ and 10^th^ days (Figures 5B, C). To validate the efficiency of the metacyclogenesis process, we used RT-qPCR to verify the transcript levels of *Metacyclin* III, a gene naturally upregulated in metacyclic forms (https://tritrypdb.org) (Figures 5D). Our results showed reduced transcript levels of *Metacyclin* III in *IP6K*^-/+^ cells compared to control during metacyclogenesis kinetics (5^th^ and 10^th^ days), confirming that this group present fewer metacyclics.

Together, these findings make it clear that the increased numbers of quiescent cells previously observed in the *IP6K*^-/+^ cells are not the result of increased numbers of metacyclic forms. Furthermore, our results suggest that low levels of IP6K impair metacyclogenesis.

### Low IP6K levels impair T. cruzi invasion and trypomastigogenesis within human cardiomyocytes

Considering that *T. cruzi* has an apparent tropism for cardiac muscle (Zhang and Tarleton, 1999; Fernandes and Andrews, 2012), we wondered whether the few metacyclic forms from *IP6K*^-/+^ cells would be capable of infecting human cardiomyocytes. To evaluate *T. cruzi* infection within human cardiomyocytes, we first stimulated metacyclogenesis for 10 days, as performed in the previous assay (Figure 5). Next, we used LLC-MK2 cells for Tissue-culture cell-derived trypomastigote forms (TCTs) multiplication and maintenance. Then, we infected human cardiomyocytes (AC16 cells) using a multiplicity of infection (MOI) 10:1 and collected samples 3, 24, 48, 72, 96 and 120 h after infection, timings expected to encompass invasion (3 h), multiplication (from 24 – 96 h) and egress (120 h) steps of the intracellular development of *T. cruzi* (Figure 6A).

**Figure 6.**
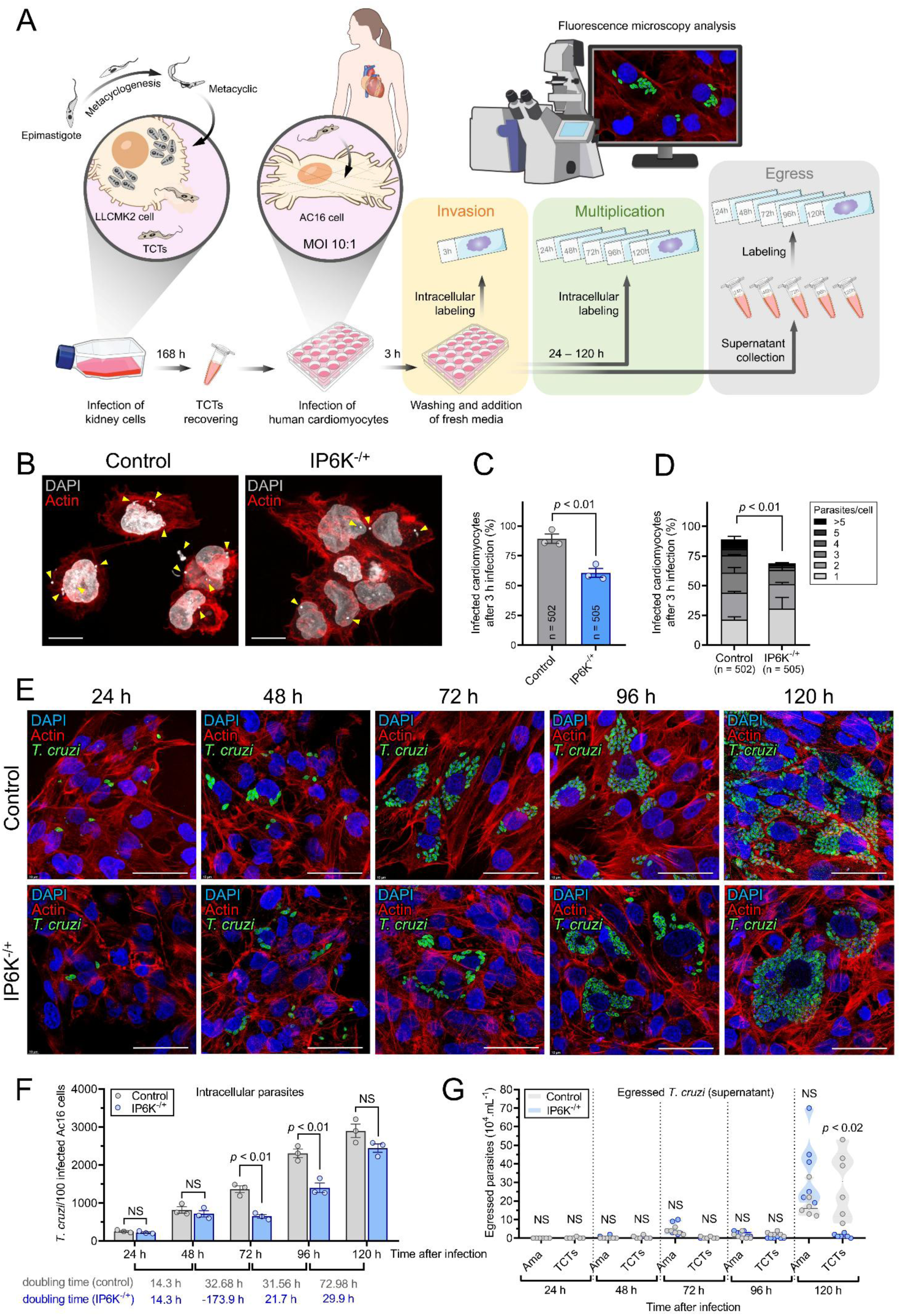
*T. cruzi* IP6K-deficient cells showed impaired invasion rate and deficient trypomastigogenesis, with an intense egress of amastigotes and few egresses of TCTs 120 h after infection within human cardiomyocytes. **A.** Scheme summarizing the approach used. **B**. Representative images of the invasion assay highlighting human cardiomyocytes containing few *T. cruzi* metacyclics. **C–D**. The percentage of cardiomyocytes infected with *T. cruzi IP6K*^-/+^ are lower relative to control. **E**. Immunofluorescence images of human cardiomyocytes infected with *T. cruzi* 24, 48, 72, 96 and 120 h after infection. F-actin filaments (red), nuclei (blue), and *T. cruzi* amastigotes (green) are shown. Bars = 50 μm. **F**. The proliferation rate of *T. cruzi IP6K*^-/+^ amastigotes is slower (relative to control) in the first 96 h but increases from 96 to 120 h. **G**. *IP6K*^-/+^ cells present intense egress of amastigotes and few egresses of TCTs 120 h after infection. The error bars in the graphs C, D and F represent the standard error of the mean (± SEM) from an assay performed in triplicate. The egress measurement was performed in hexaplicate.

After 3 h (invasion step), we observed that *IP6K*^-/+^ cells infected less cardiomyocytes relative to control (Figure 6B, C). Moreover, cardiomyocytes infected with *IP6K*^-/+^ cells showed few *T. cruzi* amastigotes per cell relative to control (Figure 6D), suggesting that lowered *IP6K* levels impair *T. cruzi* invasion within human cardiomyocytes. Next, across 24 – 120 h, when we expect to see amastigote multiplication, the number of amastigotes per infected cardiomyocyte increased consistently in the control group (Figure 6F – gray bars). In contrast, *IP6K*^-/+^ cells showed lower numbers than control at 24 and 48 h (Figure 6F – blue bars), but infection density reached that of control at 120 h (Figure 6E and F). Nevertheless, from 72 – 120 h after infection, *IP6K*^-/+^ cells showed accelerated intracellular multiplication (Figure 6F – lower doubling time values relative to control), with the average number of parasites per infected cardiomyocytes reaching the control group after 120 h (2446 ± 148 for *IP6K*^-/+^ and 2899 ± 205,1 for control).

When we look at the egressed parasites, we observed that there was no significant release from 24 – 96 h after infection for both groups control and *IP6K*^-/+^, considering amastigotes and TCTs (Figure 6G). After 120 h after infection, we observed that *IP6K*^-/+^ cells egressed from cardiomyocytes as amastigotes and only a few parasites were able to complete trypomastigogesesis and egress as TCTs (Figure 6G – 120 h). This finding suggests that IP6K-deficient amastigotes show an impaired ability to perform trypomastigogenesis, i.e., most of the intracellular population of *T. cruzi* (amastigotes) do not transform into trypomastigotes but still multiply and egress from the host cell as amastigotes.

We then decided to investigate further what might be happening 120 h after infection, which encompass trypomastigogenesis (ama–trypo transition) (Supplementary Figure 3A). First, we lysed the infected cardiomyocytes to assess the morphology of the intracellular parasites. The control group presented a similar number of amastigote and epimastigote-like forms per 100 cardiomyocytes lysed (79.3 ±2.8 and 83 ±4, respectively), a smaller number of intermediate forms (35.6 ±0.4) and only a few TCTs (6.3 ±1.5) (Supplementary Figure 3B – gray bars). This result is within expectations, since most of the TCTs were egressed from the cardiomyocytes 120 h after infection. However, when compared to the control, *IP6K*^-/+^ cells presented a significantly higher number of amastigotes (155.6 ±10.8) and lower number of epimastigote-like forms (37 ±5.3), intermediate forms (8 ±2) and TCTs (3 ±1.3) (Supplementary Figure 3B – blue bars).

This finding suggests that the observed low number of TCTs and intermediate forms in the *IP6K*^-/+^ cells is due to problems during trypomastigogenesis, since we did not observe egressed TCTs 120 h after infection (Figure 6G). In other words, the increased number of IP6K-deficient amastigotes within the human cardiomyocytes 120 h after infection suggests that IP6K contributes to trypomastigogenesis. Of note, *T. cruzi* epimastigote-like forms observed in infected cardiomyocytes correspond to intracellular transitional forms occurring during trypomastigogenesis (Almeida-de-Faria et al., 1999) and are not related to the replicative epimastigotes found *in vivo* (in the insect vector) or *in vitro* (cultivated in laboratory).

To evaluate the commitment of these different *T. cruzi* forms with the cell cycle, we subjected them to EdU incorporation for 18 h. We observed that both groups (control and *IP6K*^-/+^) presented amastigotes and epimastigote-like forms with similar proportions of EdU-labeled cells (Supplementary Figure 3C). However, the intermediate forms from *IP6K*^-/+^ cells showed a low number of EdU-labeled cells (69 ±7.3%) relative to the control (96.6 ±2.2%), which suggests a higher proportion of quiescent cells (Supplementary Figure 3C). As expected, there was no significant EdU incorporation in the TCTs forms since they are naturally quiescent (non-replicative).

Furthermore, we also tested the quiescence and replication capacity of *T. cruzi* egressed forms 120 h after infection by incorporating EdU for 18 h or 1 h, respectively. We observed that there were no significant differences regarding quiescence (Supplementary Figure 3D). However, we observed that *IP6K*^-/+^ cells presented a slight increase in EdU-labeled cells (25 ± 3.3%) compared to the control (13.5 ± 2.3%), suggesting a higher proportion of cells with replication capacity, most of them amastigotes (Supplementary Figure 3E). This result corroborates what we observed previously: a large proportion of amastigote forms are egressed 120 h after infection in the *IP6K*^-/+^ group, suggesting a defect in intracellular trypomastigogenesis.

To gain insight into potential pathways that could be altered due to the IP6K reduced levels, consequently, affecting *T. cruzi* life cycle (metacyclogenesis and trypomastigogenesis), we decided to perform transcriptomics analyses of the epimastigotes and TCTs forms.

### The differential expression of membrane and cell surface genes in response to IP6K reduced levels helps to explain the observed discrepancies during metacyclogenesis and trypomastigogenesis

We next investigated which genes are differentially regulated in response to lowered IP6K levels, aiming to identify altered pathways that might explain the observed discrepancies related to metacyclogenesis and trypomastigotegenesis. To this end, we performed transcriptomic analyses of the *T. cruzi* epimastigote and TCT forms (egressed 120 h after infection), comparing control and *IP6K*^-/+^ cells.

To assess the separation between replicates from each group (epimastigotes and TCTs), we performed a principal component analysis (PCA) using the prcomp function (Supplementary Figure 4A, D). Although there was some variance, the values among replicates were relatively close, suggesting robustness in the analyses (Supplementary Figure 4D). Moreover, to assess the quality of the reads obtained, we also quantified and mapped all reads of the samples analyzed (Supplementary Figures 4B-C). It should be noted that, due to the low number of TCTs egressed 120 h after infection in the *IP6K*^-/+^ cells (Figure 6G), we performed dozens of assays to obtain a total number of TCTs that was sufficient to compose a pool for a single transcriptomic analysis. Due to that, the percentage of quantified and mapped reads for this group was the lowest (Supplementary Figures 4B-C).

Of note, IP6K expression levels were, as expected, reduced in *IP6K*^-/+^ epimastigotes and TCTs, confirming the transcriptomics assay and analysis worked as expected.

For the epimatigotes, when comparing the control with *IP6K*^-/+^, we found 450 differentially expressed genes (DEGs) (251 upregulated and 199 downregulated) (Figure 7A – left; Supplementary Table 7). Among these genes, the most downregulated were TcCLB.508737.10 and TcCLB.508743.10, and the most upregulated were TcCLB.511913.9 and TcCLB.511367.20 (Figure 7A – left; Supplementary Table 7). TcCLB.508737.10 and TcCLB.508743.10 genes encode proteins from the TASV-C family, which are normally linked to the surface of *T. cruzi* by a GPI anchor and are spontaneously released into the medium (Bernabó et al., 2013). TcCLB.511367.20 is a hypothetical uncharacterized protein, but TcCLB.511913.9 encodes a ribonuclease L inhibitor, which is a protein that belongs to the ABC (ATP-binding cassette) family of proteins in mammals. However, in *T. cruzi* its function remains uncharacterized.

**Figure 7.**
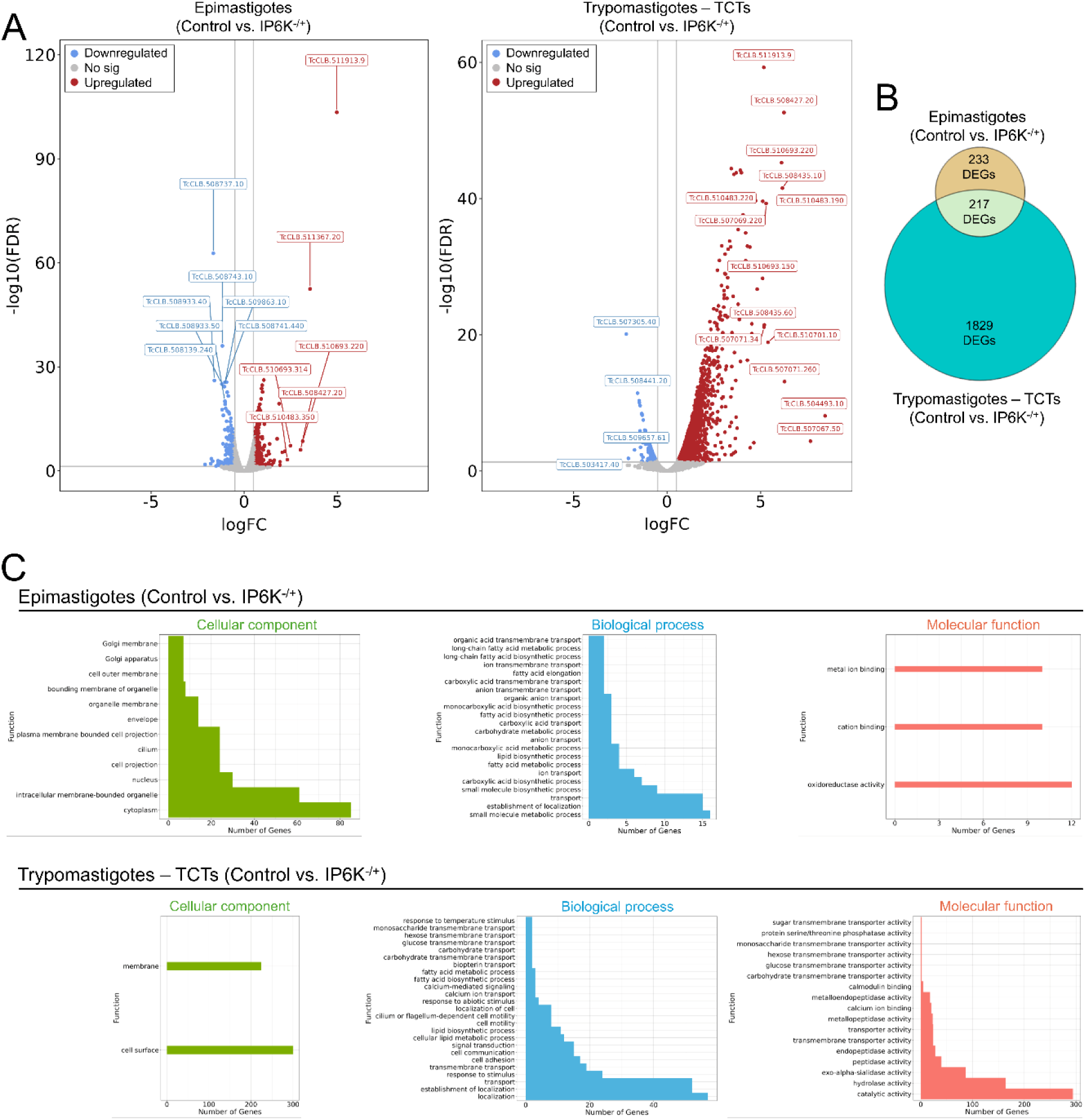
Transcriptomics analysis reveals differential expressions of membrane and cell surface genes in response to the loss of a single *IP6K* allele. **A.** Volcano plots showing DEGs in both groups analyzed epimastigotes and TCTs. **B**. Venn plot showing that 217 DEGs were differentially expressed in both groups epimastigotes and TCTs. **C**. Functional enrichment analysis using topGO and gene ontology (GO) for epimastigotes (top) and TCTs (bottom).

For the TCTs, when comparing the control with *IP6K*^-/+^, we found 2,046 DEGs (1,633 upregulated and 413 downregulated) (Figure 7A – right; Supplementary Table 8). Among these genes, the most downregulated were TcCLB.507305.40 and TcCLB.508441.20, and the most upregulated were TcCLB.511913.9 and TcCLB.508427.20 (Figure 7A – left; Supplementary Table 8). TcCLB.507305.40 gene encodes a CCCH zinc finger protein (ZC3H11). One of the homologs of this protein family has already been characterized as an essential positive regulator of *T. cruzi* differentiation from epimastigote to metacyclic forms (Alcantara et al., 2018). TcCLB.508441.20 encodes a phosphoenolpyruvate carboxykinase (PEPCK) protein, which is associated with carbohydrate catabolism in *T. cruzi* (do Amaral et al., 2021). Notably, TcCLB.511913.9 was the most upregulated gene in both epimastigotes and TCTs, although its function remains elusive in *T. cruzi*. TcCLB.508427.20 encodes a Mucin-Associated Surface Protein (MASP). MASP glycoprotein family is the second largest multigene family in *T. cruzi* and most studies indicate that MASP are surface proteins that can undergo glycosylation (Espinoza et al., 2023).

Among all DEGs identified, 217 were differentially expressed in both epimastigotes and TCTs groups (Figure 7B; Supplementary Tables 7-8). We therefore made a functional enrichment analysis using topGO (Alexa and Rahnenfuhrer, 2024) and gene ontology (GO), grouping genes into the three categories: cellular components, biological processes and molecular function (Figure 7C; Supplementary Tables 9-10). Compared to the TCT group, epimastigotes presented a higher number of DEGs related to cellular components, but on the other hand presented a lower number of DEGs related to molecular function (Figure 7C; Supplementary Tables 9-10). Furthermore, the 3 terms related to molecular function that presented DEGs in the epimastigotes (metal ion binding, cation binding and oxidoreductase activity) did not present DEGs in the TCTs. Interestingly, the only 2 terms related to cellular component that presented DEGs in the TCTs (membrane and cell surface), did not present DEGs in epimastigotes (Figure 7C; Supplementary Tables 9-10).

Although speculative, the high number of DEGs related to membrane and cell surface in the TCTs (Supplementary Table 10) helps to explain the observed discrepancies during trypomastigogenesis in response to reduced IP6K levels. The same correlation can be inferred for the epimastigotes, which presented many DEGs related to metal ion and cation binding, and oxidoreductase activity, pathways that may explain the altered morphology and impaired metacyclogenesis in response to the reduced IP6K levels.

## Discussion

The primary function of IP6K is to convert the fully mono-phosphorylated inositol (IP_6_) into the inositol pyrophosphate IP_7_ (1-IP_7_ or 5-IP_7_). However, in the last years, a growing body of work has shown moonlighting functions for IP6Ks in several cellular processes, such as neuronal migration, cell death, autophagy, nuclear translocation, cytoskeletal remodeling, cellular migration, metabolism, gene expression, DNA repair, and fungal pathogenicity (reviewed in Heitmann and Barrow, 2023; Chakkour and Greenberg, 2024).

Although *T. cruzi* IP6K is partially conserved among eukaryotes (Figure 1A, B) and retains the PxxxDxKxG motif characteristic of IP_6_-kinases (Supplementary Figures 1A, B), the loss of a single *IP6K* allele does not significantly alter PP-IPs levels (Figure 2F, G). In the qualitative analysis of PP-IPs by PAGE, we detected slower-migrating bands of unknown nature that are abundant in our IPs extracts (Figure 2F). This finding had already been reported by Wilson et al. (2015) in human cell lines and, like us, they were also unable to determine the origin of these bands. To better ascertain the PP-IP levels in our *T. cruzi* cells, we caried out PP-IPs quantification by CE-ESI-MS (Figure 2G).

Two interesting points related to this PP-IPs analysis deserve attention. First, the 1,5-IP_8_ levels observed in Control and IP6K^-/+^ cells were extremely low (∼1 fmol.μg protein^-1^) (Figure 2G), which can be explained by the absence of PP-IP5K orthologs in the *T. cruzi* genome. This calls into question the presence of IP_8_ by the PAGE assay (identified with a question mark in Figure 2F). These findings contrasts with those obtained by Mantilla et al. (2021) and Bertolini et al. (2025), where they found 1,5-IP_8_ in *T. cruzi* (Y strain). A possible explanation for this apparent discrepancy in 1,5-IP_8_ concentration may be related to *T. cruzi* strain-specific peculiarities, since our assays were carried out using the reference strain (CL Brener). The other point that deserves attention is that we normalized the PP-IPs levels by the amount of protein extracted proportionally from each sample (Figure 2F, G), which we believe is the most appropriate approach (BináLee et al., 2025). When we normalized the PP-IPs levels by the number of cells, we observed that the PP-IPs levels in IP6K^-/+^ cells appeared to decrease (*p* = 0.0403 for 1-IP_7_, *p* = 0.00167 for 5-IP_7_ and *p* = 0.68145 for 1,5-IP_8_) (Supplementary Figure 2F). However, we do not believe this is the ideal method as it does not account for potential biases unintentionally introduced between cell collection and polyphosphate extraction (BináLee et al., 2025).

The maintenance relatively stable of the PP-IP levels in the IP6K^-/+^ cells is intriguing because it suggests that *T. cruzi* IP6K itself, and not loss of its metabolic product, is responsible for the altered phenotypes observed: cell growth impairment (Figure 2A), altered cellular morphology (Figure 2H-K), increased quiescent cells (Figures 3 and 4), and impairment of metacyclogenesis, cell invasion and trypomastigogenesis (Figures 5 – 6). Through computational analysis, it was possible to observe that *T. cruzi* IP6K exhibits a disordered N-terminal portion (0–299 aa) (Supplementary Figure 1), which may confer potential allosteric sites to this kinase. However, when analyzing the *T. cruzi* IP6K N-terminal in the InterPro platform (https://www.ebi.ac.uk/interpro/) and NCBI conserved domain search (https://www.ncbi.nlm.nih.gov/Structure/cdd/wrpsb.cgi) using the data banks CATH-Gene 3D, CDD, COG, HAMAP, KOG, NCBIFAM, Panther, Pfam, PIRSF, PRINTS, PRK, PROSITE, SMART, SFLD, SMART, SUPERFAMILY, and TIGR, we did not identify any conserved motif/domain. In-depth computational assays are necessary to determine whether *T. cruzi* IP6K indeed possesses allosteric sites and potential associated moonlighting functions. Additionally, experimental methods using purified *T. cruzi* IP6K and N-terminal truncated mutants could shed light on the role of the predicted disordered region.

Yet, according to Mantilla et al. (2021), a single *IP6K* allele deletion seems to be indeed important for *T. cruzi* cell morphology and growth. Moreover, a study performed in *Crypotococcus neoformans* revealed that the knockout of *Kcs1* (*IP6K* ortholog in fungi) showed decreased growth rates compared to wild type cells (Lev et al., 2015). The authors suggested that the phenotype observed was attributed to defects in biosynthetic pathways of key macromolecules like PP-IPs, nucleotides and fatty acids that depend on mitochondrial enzymes. However, IP6K could also play a direct role in these pathways through interaction with other proteins. Additionally, based on SEM images (Figure 2H) and on the quantification of size and internal complexity by flow cytometry (Figure 2I-K), it is possible to hypothesize that structural alterations in *T. cruzi* IP6K^-/+^ cytoskeleton may be responsible for some of the described impairments in metacyclogenesis, amastigogenesis, and trypomastigogenesis, consequently affecting infective capacity (Figures 5-6). Thus, further assays capable of measuring intracellular traffic and cytoskeleton integrity with focus on the state of organization of actin and microtubules are needed to support this hypothesis. Unfortunately, our study is not comprehensive enough to provide this information.

Regarding quiescence, a recent study demonstrated that amastigotes could reach a quiescent state (called dormancy), and this process would be responsible for allowing *T. cruzi* to escape the action of antitrypanosomatid compounds (Sánchez-Valdéz et al., 2018). More recently, other studies have shown that *T. cruzi* amastigotes indeed assume a quiescent state (G0 state) when infecting mammalian cells and this process is responsible for parasite persistence (Bustamante et al., 2020; Jayawardhana et al., 2023). This stealthy behavior appears to be essential for circumventing the action of compounds that can cause DNA damage, such as Benznidazole (Jayawardhana et al., 2023). By assuming this state of non-commitment to the cell cycle (quiescence), *T. cruzi* prevents its replication machinery from generating DSBs when attempting to transpose potential DNA structure alterations caused by DNA-binding drugs, alkylating agents or other compounds. Whether *T. cruzi* can repair these DNA lesions while dormant and if these lesions can trigger quiescence are questions that deserve further investigation. It is worth mentioning that some studies already reported that homologous recombination (HR) dependent processes could activate quiescence in response to DSBs in *T. cruzi* epimastigotes (Resende et al., 2020; Costa-Silva et al., 2021). However, to our knowledge, our findings are the first to demonstrate that quiescence can be activated in the absence of DNA damage and in response to low levels of a kinase (IP6K) (Figure 4), contrary to the prevailing hypothesis that spontaneous DNA damage would trigger cell cycle arrest, leading to quiescence (Bunkofske et al., 2025).

We show here for the first time that IP6K regulates *T. cruzi* life cycle transitions. Metacyclogenesis (epi–meta transition) was significantly impaired due to the low levels of IP6K (Figure 5). In this regard, a recent study showed that histone deacetylase 4 (TcHDAC4) is important for *T. cruzi* morphology and metacyclogenesis (Picchi-Constante et al., 2021). Curiously, in mammals, IP6K1 (one of the *IP6K* paralogs) associates with chromatin and controls histone demethylation by regulating the demethylase Jumonji domain-containing 2C (JMJD2C), which catalyzes the removal of trimethyl groups from lysines 9 and 36 of histone H3 (Burton et al., 2013). In other words, it appears that IP6K can assume functions related to the regulation of histone modifications that may influence other processes, such as morphological alterations. Whether IP6K can assume this same function in *T. cruzi* and regulate histone modifications to control metacyclogenesis is an issue that requires further investigation. Another possible explanation for the impaired metacyclogenesis observed in IP6K-deficient cells is oxidative stress management. In mammalian cells, IP6K (IP6K2 paralog) participates in oxidative stress responses through binding with creatine kinase-B (CK-B) to maintain mitochondrial function and keep ROS levels constant (Nagpal et al., 2021). However, we believe this is not occurring in our cells, since mitochondria from IP6K-deficient *T. cruzi* cells appear to function normally, even at low IP6K levels, as observed in our metabolic viability assays using MTT, which measures mitochondrial activity (Figure 4I). This is justified by the low similarity between *T. cruzi* IP6K and human IP6K2, which is ∼25% (Supplementary Figure 1). In other words, as the interaction between human IP6K2 and CK-B possibly does not involve the catalytic ArgRIII domain, as it is a moonlight function of IP6K (Nagpal et al., 2021, 2022), it is more likely that this interaction is not present in *T. cruzi* since the only conserved region between *T. cruzi* IP6K and human IP6K2 corresponds to the motifs located within the catalytic ArgRIII domain (Supplementary Figure 1).

We also observed that *T. cruzi* trypomastigogenesis (ama-trypo transition) was impaired within human cardiomyocytes in response to lowered levels of IP6K (Figure 6). A study performed 10 years ago showed that *T. cruzi* trypomastigogenesis is dependent on an intracellular Ca^2+^ channel and a key molecule involved in Ca^2+^ signaling, called inositol 1,4,5-trisphosphate receptor (TcIP3R) (Hashimoto et al., 2015). IP3R is also a major regulator of apoptotic signaling in mammals (Vicencio et al., 2009) and autophagy in *T. cruzi* (Chiurillo et al., 2020). Also, in *T. cruzi*, TcIP3R is predominantly localized in acidocalcisomes (Chiurillo et al., 2020), acidic calcium stores rich in polyphosphates presented in trypanosomes (Vercesi et al. 1994; Docampo et al., 1995). Interestingly, the PP-IPs synthesis pathway in *Trypanosoma brucei* is linked to polyphosphate synthesis in acidocalcisomes (Cordeiro et al., 2017). Based on this evidence, we hypothesize that IP6K may influence TcIP3R levels in the acidocalcisomes, regardless of PP-IPs, and this could explain the altered trypomastigogenesis observed by us.

Lowered levels of IP6K also appear to contribute to an increased replication rate of *T. cruzi* amastigotes (Figure 6F; Supplementary Figure 3E). Interestingly, although there is evidence that PP-IPs are responsible for promoting increased replication rate in cancer cells by antagonizing the action of hepatic kinase B1 in mice (Rao et al., 2014), *IP6K1* knockout mice present accelerated tumor cell growth (Lee et al., 2022). These findings, although seemingly paradoxical given that IP6K is responsible for PP-IP synthesis, can be easily explained if we assume that IP6K has moonlighting functions. In other words, the data obtained in mice regarding tumorigenic growth corroborates our findings, especially if we assume that IP6K-deficient *T. cruzi* amastigotes undergo apparent uncontrolled proliferation (Figure 6F,G), which is a primary characteristic observed in cancer cells (Lun et al., 2015). Furthermore, amastigotes with increased replication rates could contribute to sustaining an infectious cycle in cardiomyocytes (Ley et al., 1988; Stahl et al., 2013; Bonfim-Melo et al., 2018). Further studies are needed to monitor reinfection by amastigotes after egress (i.e.: after 96 or 120 h pos infection) and to confirm whether the IP6K-deficient forms could indeed infect actively more AC16 cells.

Finally, our transcriptomics analysis revealed that the single *IP6K* allele deletion contributed to DEGs associated with membrane and cell surface in trypomastigotes (TCTs) and epimastigotes (Figure 7, Supplementary Tables 7-8, Supplementary Figure 5A, B), which helps to explain the observed alterations during invasion and infection. The main DEGs found in TCTs were related to MASP family, Trans-sialidases family (Gp85, Gp82 Gp90 and Gps 35/50 proteins), Mucins family, Cruzipain family, and Oligopeptidase B, which are essential for *T. cruzi* invasion, infection, and host immune evasion (Supplementary Table 8; Supplementary Figure 5B). In general, Trans-sialidases are involved in the transfer of sialic acid from host glycoconjugates to mucins of the parasite, which are the major family of glycoproteins expressed in the cell surface in epimastigotes and trypomastigotes (Giorgi and de Lederkremer, 2011). Members from MASP family constitute parasite antigens that are recognized by the host antibodies. The large repertoire of MASP sequences contribute to the ability of *T. cruzi* to infect different cell types (dos Santos et al., 2012). Thus, when these glycoproteins are altered, *T. cruzi* fails to establish an efficient infection in the host cell, whether *in vivo* or *in vitro* (Manque et al., 2011; dos Santos et al., 2012; Leão et al., 2022). We observed that members of MASPs and gp35/50 mucins molecules are upregulated in TCTs IP6K-deficient parasites relative to control (Supplementary Figure 5B). In *T. cruzi* metacyclic forms, gp35/50 mucins mediate invasion through interaction with host cell annexin A2 and clathrin-dependent endocytosis (Onofre at al., 2022), while MASPs facilitate the adhesion of trypomastigotes to host cells, including cardiomyocytes (Espinoza et al., 2023). Thus, the upregulation of these protein families should increase invasion/infection in the IP6K^-/+^ lineage. However, this was not observed in our assays (Figure 6B-D).

The explanation for this finding may lie in the fact that our transcriptomics data also revealed that Oligopeptidase B and cruzipain family members are predominantly downregulated in IP6K-deficient parasites (Supplementary Figure 5). Oligopeptidase B is an important serine protease expressed in all developmental forms of *T. cruzi* that act as a key virulence factor during host cell invasion and infection (reviewed in Beatriz-Vermelho et al., 2010). This protease enables *T. cruzi* to enter mammalian cells by generating a soluble signaling molecule that triggers intracellular calcium transients, leading to host cell lysosome recruitment and fusion (Caler et al., 1998). Cruzipain is crucial papain-like cysteine protease, also expressed in all developmental forms of *T. cruzi* that assists the parasite in entering host cells by mobilizing bradykinin and endothelin receptors during invasion (Scharfstein et al., 2000; Andrade et al., 2012; Santos et al., 2021). Also, cruzipain helps *T. cruzi* escape the host’s adaptive immune system and it is essential for differentiation and intracellular growth (Doyle et al., 2011). Thus, the impaired invasion rate and deficient trypomastigogenesis in the *T. cruzi* IP6K^-/+^ lineage can be explained by the downregulation of Oligopeptidase B and cruzipain, which seems be critical for these processes (Duschak and Couto, 2009; Motta et al., 2012).

However, it is important to note that our study is limited in providing direct evidence to justify the observed phenotypic alterations associated with infections and differentiation processes. The observed reduction in infectivity and impairment in metacyclogenesis, amastigogenesis, and trypomastigogenesis may represent indirect or stage-specific consequences, rather than a single continuous mechanistic pathway. Thus, although challenging, further and tightly regulated inducible assays are necessary to discriminate between these two possibilities. Furthermore, it is worth mentioning we tried to rescue the original phenotype of our IP6K-deficient *T. cruzi* cells by reintroducing the IP6K allele cloned into pROCK plasmid (daRocha et al., 2004). However, the IP6K ectopic expression did not work, as we were unable to select any transfected cells. We also tried to reintroduce the IP6K allele into a constitutive ribosomal locus using pTREX plasmid (Lorenzi et al., 2003). Again, we were unable to select any transfected parasites. These failed attempts suggest that controlling endogenous IP6K levels may be overly precise in this parasite. Thus, further assays using trypanosomatids inducible gene edit systems, such as DiCre (Duncan et al., 2016; Yagoubat et al., 2020) are necessary to confirm this hypothesis.

Taken together, our findings strongly suggest that IP6K is important for normal cell growth and maintenance of the default morphology in *T. cruzi* epimastigotes. Also, IP6K seems to regulate the entrance of *T. cruzi* epimastigotes in a quiescent state, whose function and mechanism remain undefined. Furthermore, we presented here, for the first time, robust data that supports the idea that IP6K is critical to sustain *T. cruzi* life-cycle transitions (in a directly or indirectly way), possibly independently of PP-IPs, since IP_7_ levels are not altered in response to the single *IP6K* allele depletion (Figure 2F, G), as evidenced by other study (Mantilla et al., 2021). Since disruption of both *IP6K* alleles did not yield viable parasites and the similarity to its human homolog is low, this kinase represents a promising target for drug development against Chagas disease.

## Acknowledgments

The authors are grateful to Dr. Débora Andrade-Silva (The University of Texas – Southwestern Medical Center, USA) for her critical revision and valuable suggestions to improve the manuscript’s quality, and Dr. Marija Krasilnikova (The University of Glasgow, Scotland, UK) for her help with WGS analysis. We also acknowledge Marcelo Nunes Silva for his immeasurable help in the preparation of reagents/solutions and laboratory organization. Finally, we would like to thank Professors Maria Isabel N. Cano (São Paulo State University, Brazil), Bianca S. Zingales (University of São Paulo, Brazil), Maurício Baptista (University of São Paulo, Brazil), and Alexander Henning Ulrich (University of São Paulo, Brazil) for sharing their equipment, which were of paramount importance for the development of this work.

---

**Supplementary Figure 1.**
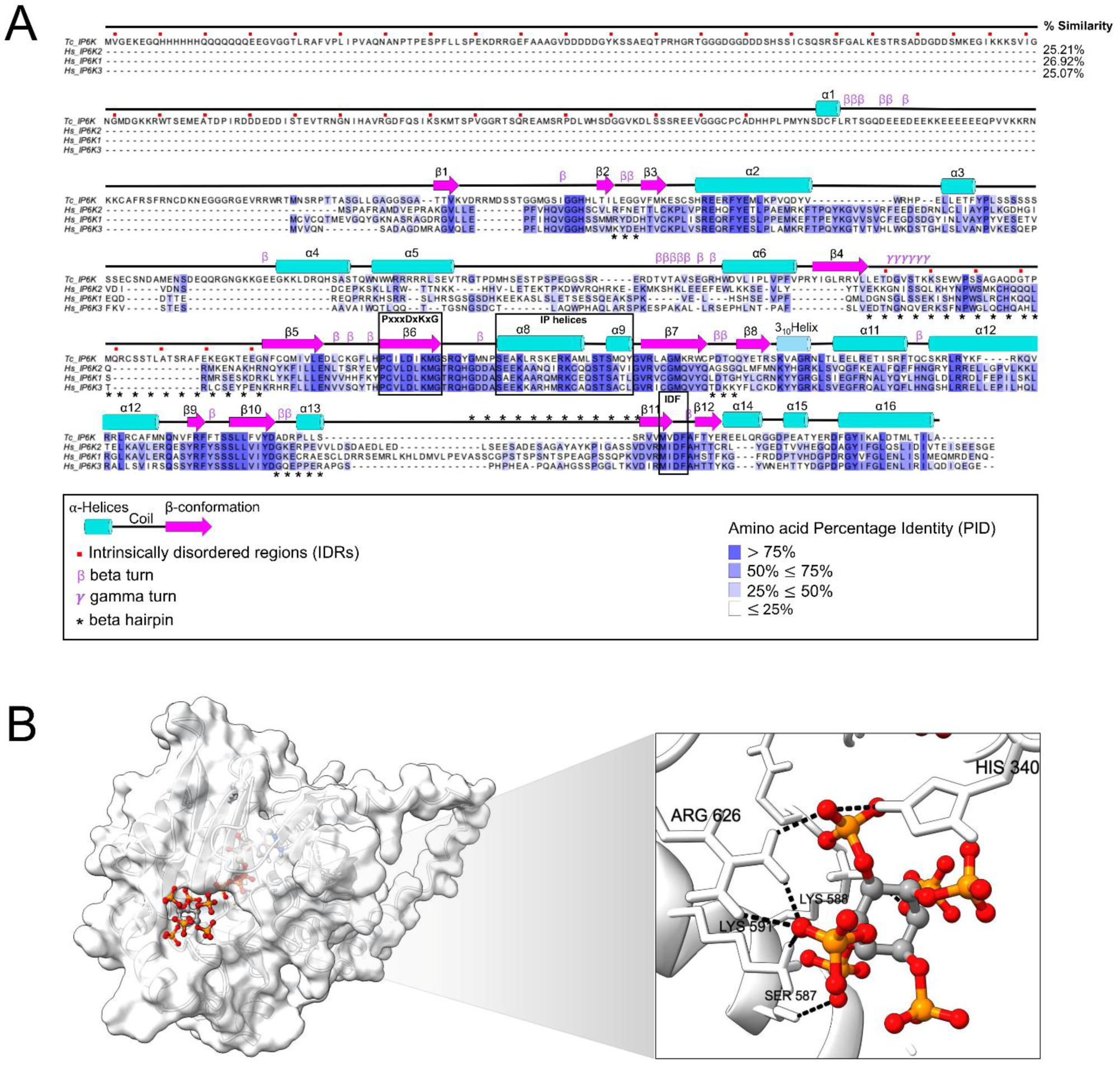
*In silico* analyses of *T. cruzi* IP6K. **A.** Alignment among *T. cruzi* IP6K and three IP6K human isoforms (*Hs*_IP6K1, *Hs*_IP6K2, and *Hs*_IP6K3). The main secondary structures are depicted according to the legend. The main motifs are highlighted with a black box. **B**. Molecular docking analyses between *T. cruzi* IP6K and IP6, the precursor of IP7. The region containing the catalytic domain is being enlarged. Important amino acid residues are highlighted and were predicted to be essential for the interaction with IP6.

**Supplementary Figure 2.**
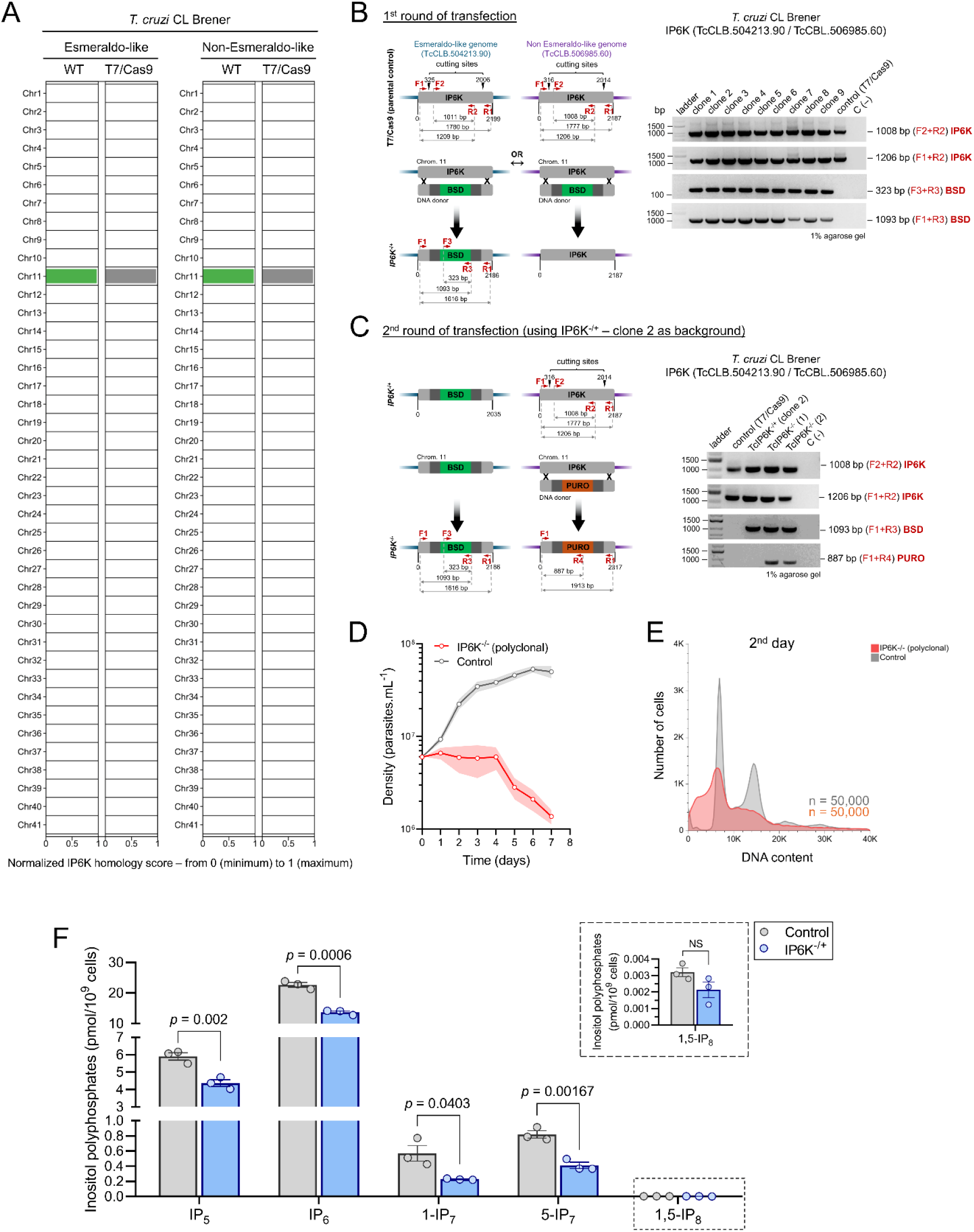
CRISPR/Cas9 strategy was used to deplete *T. cruzi IP6K*. **A.** Genome-wide distribution of IP6K homology across the *T. cruzi* CL Brener chromosomes from wild type (WT) and T7/Cas9 lineages. The plotted bar represents the normalized IP6K homology score, combining query coverage and sequence identity on a scale from 0 (minimum) to 1 (maximum). **B, C** (left). Schemes showing the approach used to deplete *IP6K* from *T. cruzi* CL Brener genome. The red arrows represent the primers designed to confirm the deletions. **B** (right). After the 1^st^ round of transfection, we only got single-allele mutants for *IP6K* (*IP6K*^-/-^). **C** (right). Using one of these *IP6K*^-/-^ clones (clone 2), we carried out a 2^nd^ round of transfection to try to obtain a *IP6K* null mutant (*IP6K*^-/-^) lineage. **D.** Growth curves showing the impaired proliferation of the polyclonal *IP6K*^-/-^ lineage. **E.** DNA content analysis showing the increased sub-G1 population of the polyclonal *IP6K*^-/-^ lineage. **F.** CE–ESI–MS analysis of polyphosphates levels (IP5, IP6, 1-IP7, 5-IP7, and 1,5-IP8) in Control and IP6K^-/+^ lineages, normalized by the number of total cells. The error bars represent the standard error of the mean (± SEM) from an assay performed in triplicate. NS means not significant; the *p*-values were obtained through Student’s t-test.

**Supplementary Figure 3.**
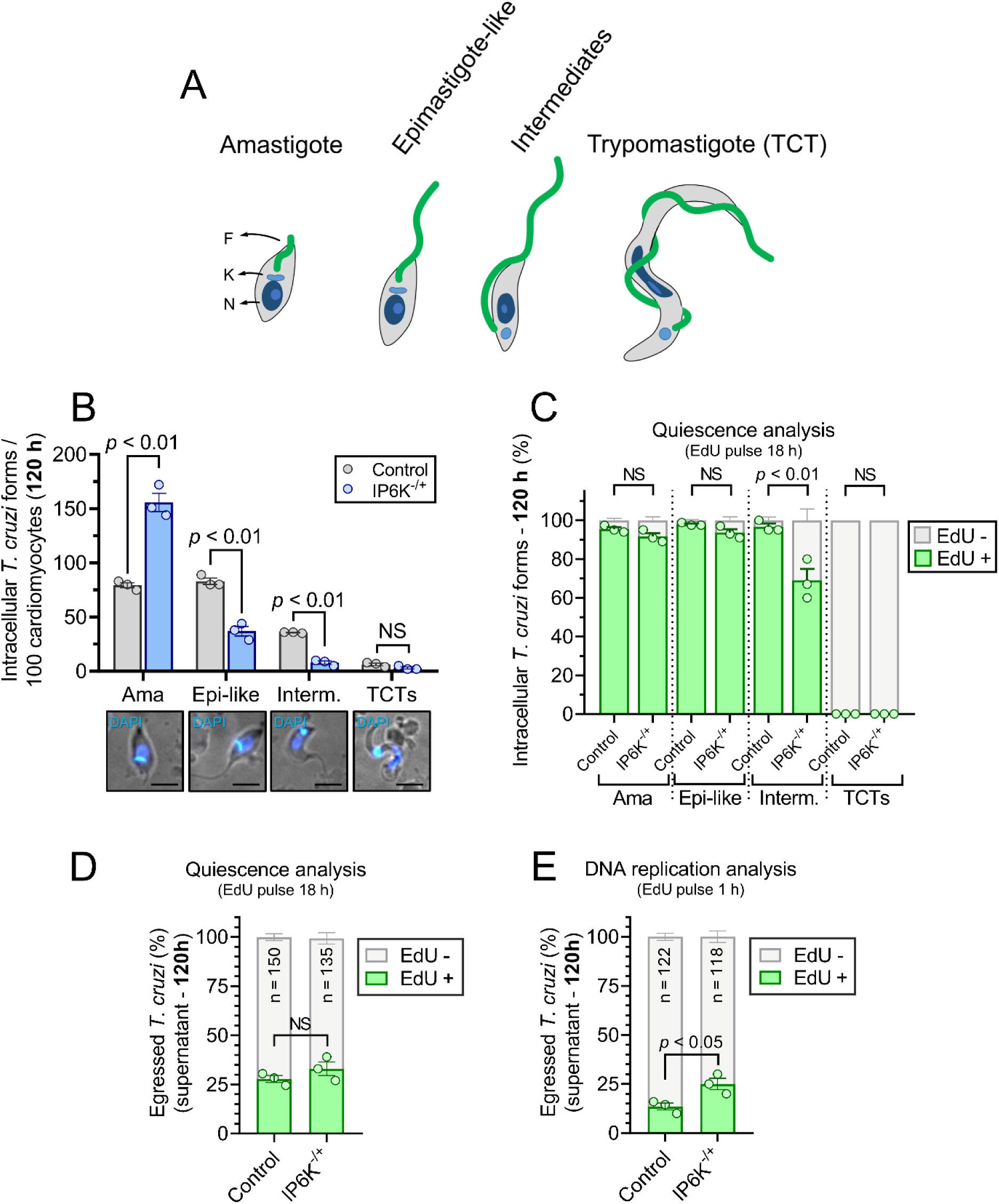
*IP6K*^-/+^ cells present part of the intracellular intermediates in a quiescent state and the egressed amastigotes have high replication capacity. **A**. Schematic representation of the main *T. cruzi* life forms during trypomastigogenesis. **B**. After 120 h, *IP6K*^-/+^ cells show more amastigotes but less epimastigote-like and intermediate forms within human cardiomyocytes. **C**. Curiously, after 120 h, IP6K^-/+^ shows a significant high number of intermediate *T. cruzi* forms in a quiescence-like (dormant) state within human cardiomyocytes. **D–E**. After 120 h, the released *T. cruzi* amastigote forms did not present differences relative to quiescence between *IP6K*^-/+^ and control (D). However, *IP6K*^-/+^ cells show more replicating cells (E).

**Supplementary Figure 4.**
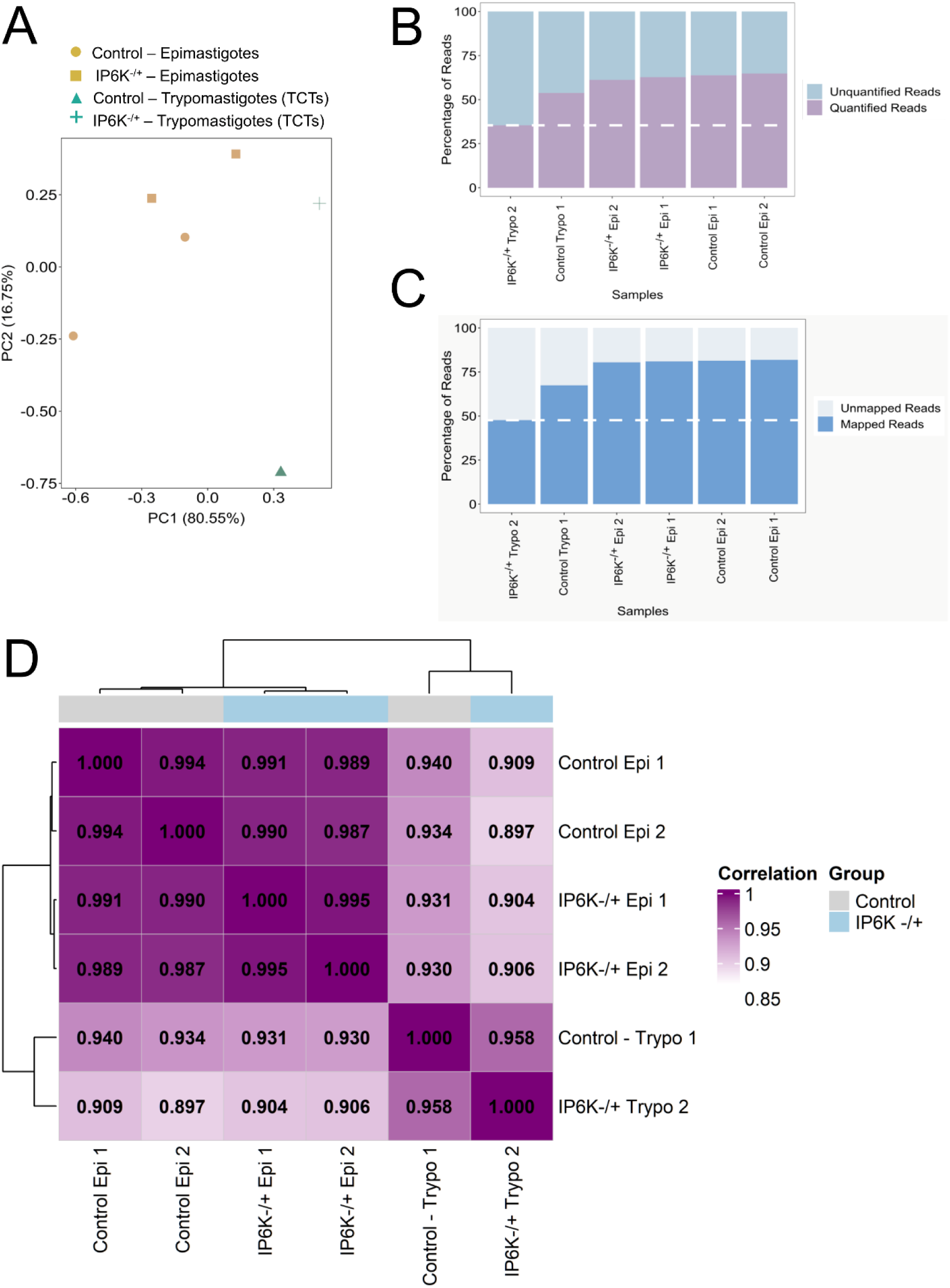
Principal component and RNA-seq reads analysis. **A**. Principal component analysis (PCA) using the prcomp function. **B.** RNA-seq reads quantification for the analyzed groups. **C.** RNA-seq reads mapping for the analyzed groups. **D**. Pairwise correlation analyses across all samples.

**Supplementary Figure 5.**
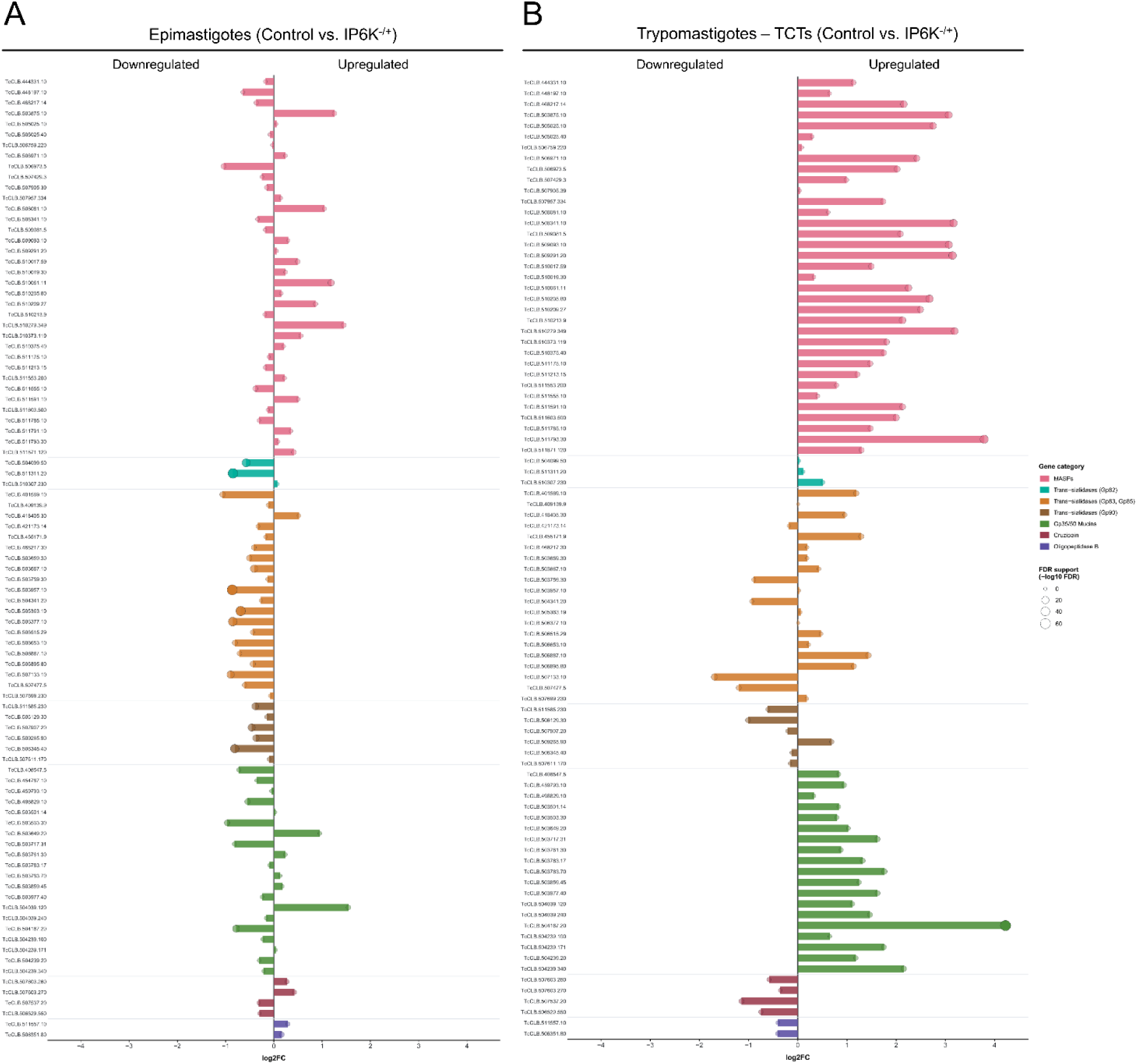
Analysis of differentially expressed virulence-related genes in response to low levels of IP6K. **A** and **B**. Differentially expressed virulence-related genes (MASPs, Trans-sialidases, Mucins, Cruzipain and Oligopeptidase B families) in both groups analyzed: *T. cruzi* epimastigotes (shown in A) and *T. cruzi* TCTs (shown in B). The representative gene set for each family (MASPs, Trans-sialidases, Mucins, Cruzipain and Oligopeptidase B families) was chosen randomly in https://tritrypdb.org.

