## Supplementary tables 1-6 for "Reduced levels of inositol hexakisphosphate kinase (IP6K) impair life-cycle transitions and the intracellular development of *Trypanosoma cruzi* within human cardiomyocytes"

#### Supplementary tables 1 – 6 from the manuscript entitled:

### Reduced levels of inositol hexakisphosphate kinase (IP6K) impair life-cycle transitions and the intracellular development of *Trypanosoma cruzi* within human cardiomyocytes

Bryan E. Abuchery[1,2,#], Thaise L. Teixeira[1,#], Vitor L. da Silva[1,2], Rauni B. Marques[3], Fernanda M. Gerolamo[1], Carolina M. C. Catta-Preta[4], Bruno A. Santarossa[5], Craig Lapsley[6], Yete G. Ferri[2], Maria Cristina M. Motta[7], Samuel C. Teixeira[8], Miguel A. Chiurrilo[9], Noelia M. Lander[9], Simone G. Calderano[5], Eduardo M. Reis[1], Richard McCulloch[6], Marcelo S. da Silva[1,2,*]

[1]Department of Biochemistry, Institute of Chemistry, University of São Paulo (USP), São Paulo, SP, Brazil; [2]Department of Genetics, Microbiology and Immunology, Biosciences Institute, São Paulo State University (UNESP), Botucatu, São Paulo, Brazil; [3]Interunit Graduate Program in Bioinformatics, University of São Paulo, SP, Brazil; [4]Department of Parasitology, Institute of Biomedical Sciences, University of São Paulo, São Paulo, SP, Brazil; [5]Butantan Institute, São Paulo, SP, Brazil; [6]The University of Glasgow, Centre for Parasitology, the Wellcome Centre for Integrative Parasitology, University of Glasgow, School of Infection and Immunity, Glasgow, UK; [7]Institute of Biophysics Carlos Chagas Filho, Federal University of Rio de Janeiro, Rio de Janeiro, RJ, Brazil, [8]Department of Immunology, Institute of Biomedical Science, Federal University of Uberlândia, Uberlândia, MG, Brazil; [9]Department of Biological Sciences, University of Cincinnati, Cincinnati, OH, USA.

**Supplementary Table 1.** Primers used for *T. cruzi* CL Brener genome editing and checking

| Primer name | Sequence (5' – 3') | Function |
|---|---|---|
| Upstream foward | TGGTGATGATGATAGTCATAGTAGTATTTGGTATAATG CAGACCTGCTGC | DNA donor amplification |
| Downstream reverse | GCAGCGGGCGATCGGCGTCGTACACAAACACCAATTT GAGAGACCTGTGC | DNA donor amplification |
| Upstream 'sgRNA' | GAAATTAATACGACTCACTATAGGATAGCCAGAGCAGG AGCTTTGGTTTTAGAGCTAGAAATAGC | Upstream 'sgRNA' syntesis |
| Downstream 'sgRNA' | GAAATTAATACGACTCACTATAGGGCAACGAGGACGTG AAAAAGGTTTTAGAGCTAGAAATAGC | Downstream 'sgRNA' syntesis |
| Common 'sgRNA' | AAAAGCACCGACTCGGTGCCACTTTTTCAAGTTGATAACG GACTAGCCTTATTTTAACTTGCTATTTCTAGCTCTAAAAC | Common primer for 'sgRNA' syntesis |
| F1 | CGATGGTGGTGATGATGATAGTC | Checking for the presence of IP6K gene |
| R1 | GCGATCGGCGTCGTACAC | Checking for the presence of IP6K gene |
| F2 | GGACGACGATGAGGATGACATTAG | Checking for the presence of IP6K gene |
| R2 | CCAATGTATCGCGGAACGAAC | Checking for the presence of IP6K gene |
| F3 | CAGCATCCCCATCTCTGAAGAC | Checking for the presence of blasticidin resistance gene |
| R3 | CATAACCAGAGGGCAGCAATTC | Checking for the presence of blasticidin resistance gene |
| R4 | GACTGACGCCAACTGTCG | Checking for the presence of puromycin resistance gene |

**Supplementary Table 2.** Primers used for RT-qPCR analyses

| Primer name | Sequence (5' – 3') | Target |
|---|---|---|
| GAPDH Fw | AGCGCGCGTCTAAGACTTACA | GAPDH (TcCLB.503687.20) |
| GAPDH Rv | TGGAGCTGCGGTTGTCATT | GAPDH (TcCLB.503687.20) |
| IP6K Fw | AACGACGGTGAAGGTTGATAG | IP6K (TcCLB.504213.90 and TcCLB.506985.60) |
| IP6K Rv | CCACCCTCCAGAATGGTTAAG | IP6K (TcCLB.504213.90 and TcCLB.506985.60) |
| MetIII Fw | ACGGTACGCTCTCTCCTTT | Metacyclin III (TcCLB.510943.44) |
| MetIII Fw | CATGCTGTCATCTTCGTTATCC | Metacyclin III (TcCLB.510943.44) |

**Supplementary Table 3**. Instrument and scan source parameters of QQQ.

| Parameter | Setting |
|---|---|
| Gas temperature | 150 °C |
| Gas flow | 11 $L.min^{-1}$ |
| Nebulizer pressure | 8 psi |
| Sheath gas temperature | 175 °C |
| Sheath gas flow | 8 $L.min^{-1}$ |
| Capillary voltage | −2000 V |
| Nozzle voltage | 2000 V |
| High-pressure RF (ion funnel) | 70 V |
| Low-pressure RF (ion funnel) | 40 V |

**Supplementary Table 4.** MRM transitions setting of inositol phosphates for samples.

| Molecular name | Precursor ion (m/z) | Product ion (m/z) | Dwell (ms) | Fragmentor (V) | Collision energy (V) | Cell accelerator (V) | Polarity |
|---|---|---|---|---|---|---|---|
| $InsP_5$ | 579.0 | 498.9 | 50 | 166 | 9 | 3 | Negative |
| $^{13}C_6$-$InsP_5$ | 585.0 | 504.9 | 50 | 166 | 9 | 3 | Negative |
| $InsP_6$ | 659.0 | 480.9 | 50 | 166 | 13 | 4 | Negative |
| $^{13}C_6$-$InsP_6$ | 665.0 | 486.9 | 50 | 166 | 13 | 4 | Negative |
| PP-$InsP_5$ | 739.0 | 319.9 | 50 | 166 | 9 | 3 | Negative |
| $^{13}C_6$-5-PP-$InsP_5$ | 745.0 | 322.9 | 50 | 166 | 9 | 3 | Negative |
| $(PP)_2$-$InsP_4$ | 819.0 | 359.8 | 50 | 166 | 9 | 1 | Negative |
| $^{13}C_6$-$(PP)_2$-$InsP_4$ | 825.0 | 362.8 | 50 | 166 | 9 | 1 | Negative |

**Supplementary Table 5.** Isotopic internal standards and final working concentrations for quantitative CE–MS analysis

| Isotopic internal standard | Inositol phosphate species | Final [ ] in sample (μM) |
|---|---|---|
| $^{13}C_6$-Ins(1,3,4,5,6)$P_5$ | Inositol pentakisphosphate ($IP_5$) | 4 |
| $^{13}C_6$-$InsP_6$ | Inositol hexakisphosphate ($IP_6$) | 8 |
| $^{13}C_6$-5-PP-$InsP_5$ | 5-diphospho-inositol pentakisphosphate (5-$IP_7$) | 1 |
| $^{13}C_6$-1- PP-$InsP_5$ | 1-diphospho-inositol pentakisphosphate (1-$IP_7$) | 1 |
| $^{13}C_6$-1,5-$InsP_8$ | 1,5-bis-diphospho-inositol tetrakisphosphate (1,5-$IP_8$) | 0.5 |

**Supplementary Table 6.** Final deletion candidates after all filters (20 kb and 1 kb analysis). Per locus summary including chromosome, coordinates, analysis scale, mean re-centered log2 depth in IP6K$^{-/+}$ and T7/Cas9, differential amplitude (ΔLog2 = IP6K$^{-/+}$ – T7/Cas9), fraction of reads with mapping quality zero (MAPQ0), split-read counts at interval edges, and final PASS status.

| Chrom | Start | End | Scale | Mean log2 IP6K | Mean log2 T7/Cas9 | Δ (IP6K – T7/Cas9) | MAPQ0 | Split reads | PASS |
|---|---|---|---|---|---|---|---|---|---|
| TcChr11-S | 261000 | 262000 | 1 kb | -inf | 1.00 | 0 | 1.00 | 0 | No |
| TcChr19-S | 1660589 | 1663589 | 1 kb | -1.105 | -1.007 | -0.098 | 1.00 | 0 | No |
| TcChr24-S | 1696100 | 1697100 | 1 kb | -1.096 | 0.953 | -0.143 | 1.00 | 0 | No |
| TcChr24-S | 1700100 | 1703676 | 1 kb | -1.047 | -1.147 | 0.100 | 1.00 | 0 | No |
| TcChr24-S | 1723776 | 1724776 | 1 kb | -1.047 | -0.980 | -0.066 | 1.00 | 0 | No |
| TcChr24-S | 1735776 | 1737776 | 1 kb | -1.137 | -0.909 | -0.228 | 1.00 | 0 | No |
| TcChr30-S | 1197998 | 1198998 | 1 kb | -3.575 | -0.332 | -3.243 | 1.00 | 0 | No |
| TcChr31-S | 445270 | 494600 | 20 kb | -0.821 | 0.077 | -0.898 | 1.00 | 0 | No |
| TcChr31-S | 1218014 | 1223014 | 1 kb | -1.246 | 0.424 | -1.670 | 1.00 | 0 | No |
| TcChr31-S | 1224014 | 1227014 | 1 kb | -1.177 | 1.079 | -2.256 | 1.00 | 0 | No |
| TcChr31-S | 1228014 | 1234014 | 1 kb | -1.191 | 0.985 | -2.176 | 1.00 | 0 | No |
| TcChr31-S | 1236014 | 1495736 | 20 kb | -0.937 | -0.045 | -0.892 | 1.00 | | Yes |
| TcChr31-S | 1247014 | 1249014 | 1 kb | -1.067 | -0.072 | -0.995 | 1.00 | 0 | No |
| TcChr31-S | 1288014 | 1291014 | 1 kb | -1.122 | -0.165 | -0.957 | 1.00 | 0 | No |
| TcChr31-S | 1294014 | 1298014 | 1 kb | -1.038 | -0.297 | -0.740 | 1.00 | 0 | No |
| TcChr31-S | 1314014 | 1316014 | 1 kb | -1.249 | -0.042 | -1.208 | 1.00 | 0 | No |
| TcChr31-S | 1332014 | 1335014 | 1 kb | -1.163 | -0.175 | -0.988 | 1.00 | 0 | No |
| TcChr31-S | 1378014 | 1381014 | 1 kb | -1.112 | -0.310 | -0.802 | 1.00 | 0 | No |
| TcChr31-S | 1423014 | 1427014 | 1 kb | -1.141 | -0.109 | -1.032 | 1.00 | 0 | No |
| TcChr31-S | 2065997 | 2146728 | 20 kb | -0.921 | -0.094 | -0.827 | 1.00 | | Yes |
| TcChr31-S | 2087997 | 2090997 | 1 kb | -1.026 | -0.139 | -0.887 | 1.00 | 0 | No |
| TcChr31-S | 2145997 | 2152997 | 1 kb | -1.095 | -0.202 | -0.893 | 1.00 | 0 | No |
| TcChr31-S | 3350610 | 3353610 | 1 kb | -1.110 | -0.184 | -0.926 | 1.00 | 0 | No |
| TcChr31-S | 3378610 | 3381610 | 1 kb | -1.131 | -0.207 | -0.923 | 1.00 | 0 | No |
| TcChr7-S | 116052 | 119052 | 1 kb | -1.114 | -0.838 | -0.276 | 1.00 | 0 | No |
